# Translocation-driven chr19-der(14) interaction is associated with disease-specific transcriptional programs in mantle cell lymphoma

**DOI:** 10.64898/2026.06.05.730311

**Authors:** Anna Schwager, Yelena Grishina, Ivan Tsimailo, Gyunel Rashidova, Eugenia Tiukacheva, Mariia Pestriakova, Lisa Lagorgette, Daria A. Bogdanova, Azat Garaev, Jean-Marie Michot, Vincent Ribrag, Oleg Demidov, Guillaume Andrieu, Sergei V. Ulianov, Sergei V. Razin, Diego Germini, Yegor Vassetzky

## Abstract

Mantle cell lymphoma (MCL) is defined by the t(11;14)(q13;q32) translocation, which drives constitutive *CCND1* expression, yet the broader regulatory consequences of this rearrangement remain incompletely understood. Here, we combined transcriptomic, epigenomic, Hi-C, and 3D-FISH analyses in primary MCL samples and cell lines to investigate the genome-wide impact of t(11;14) on chromatin organization and gene regulation beyond the rearranged chromosomes. We show that MCL cells exhibit widespread enhancer activation, accompanied by expansion of super-enhancer regions. The translocated *CCND1* locus repositions toward the nuclear interior and acquires enhancer-like features. Genome-wide analysis highlighted chromosome 19 as a hotspot of transcriptional upregulation. Notably, Hi-C and 3D-FISH revealed recurrent interchromosomal contacts between chromosome 19 and the *CCND1* locus, preferentially involving the derivative chromosome. These contacts colocalize with active RNA polymerase II and are associated with increased expression of nearby chr19 genes, suggesting an association between spatial proximity and transcriptional activation in trans. Minnelide treatment reduced chromatin accessibility at the *CCND1* locus, decreased the frequency of chr19-der14 interactions, and partially reversed the MCL transcriptional program, while exerting strong anti-tumor effects *in vitro* and *in vivo*. Together, these findings identify a recurrent interchromosomal interaction associated with coordinated gene activation in MCL and suggest that spatial genome reorganization contributes to disease-specific transcriptional programs.

## INTRODUCTION

Mantle cell lymphoma (MCL) is an aggressive mature B-cell malignancy, with a generally unfavorable prognosis, particularly in relapsed cases [1,2]. The genetic hallmark of MCL is the t(11;14)(q13;q32) chromosomal translocation, detected in over 95% of cases. T(11;14) juxtaposes the *CCND1* gene to a strong enhancer within the immunoglobulin heavy chain (IGH) locus and results in constitutive *CCND1* expression [3,4]. However, experimental studies demonstrate that *CCND1* overexpression alone is insufficient to induce malignant transformation [5,6], suggesting that additional regulatory mechanisms contribute to MCL pathogenesis.

Chromosomal translocations are increasingly recognized to exert effects beyond local enhancer hijacking, including alterations in higher-order chromatin organization and nuclear architecture. Genome organization plays a critical role in gene regulation by constraining enhancer-promoter communication within three-dimensional nuclear space [7–9]. In this context, spatial repositioning of genomic loci, e.g. following chromosomal translocation, can influence transcriptional activity by altering their proximity to transcriptionally permissive nuclear compartments and regulatory hubs [10–12]. Indeed, in MCL, the translocated *CCND1* allele was reported to relocate closer to the nucleolus, to the vicinity of active Pol II sites [13], and the major MCL translocation breakpoint corresponds to an open chromatin region decorated with active histone marks (H3K4me1/H3K4me3/H3K27ac) [14]. MCL cells were shown to undergo disease-specific chromatin compartment changes, with the tendency towards compartment activation [15], potentially favoring the formation of active transcription hubs. A recent study demonstrated that MCL-associated translocation could induce coordinated transcriptional changes of multiple genes covering entire chromosome arms, transforming these genomic segments into large regulatory units [16]. Such regulons might arise from both short-range and long-range chromatin interactions, potentially including interchromosomal contacts. Consistent with this notion, reorganization of the interchromosomal interaction landscape, particularly involving chromosomes 11 and 14, has been reported in MCL [16].

Despite these advances, the extent to which specific genomic regions at the translocation breakpoint participate in defined spatial interactions, and how such interactions influence transcriptional outputs beyond the rearranged chromosomes (chr11 and chr14), remains unclear. In particular, it is not known whether breakpoint-associated regions can act as focal points for recurrent long-range contacts that extend to other chromosomes, and whether such interactions are linked to coordinated gene expression changes outside the translocation partners. Addressing these questions is important in order to understand how chromosomal translocations can drive genome-wide transcriptional deregulation beyond local effects and to identify potential regulatory dependencies that could be therapeutically targeted.

Here, we combined transcriptomic, epigenomic, Hi-C, and 3D-FISH analyses to investigate the consequences of t(11;14) in MCL, with a focus on genome-wide transcriptional, epigenetic and spatial changes. We show that MCL cells exhibit widespread activation of the enhancer landscape and that the translocated *CCND1* locus repositions toward the nuclear interior, where it acquires enhancer characteristics. We further identify chromosome 19 as a hotspot of transcriptional activation in MCL and demonstrate recurrent, transcriptionally active interchromosomal contacts between chromosome 19 and the translocated *CCND1* locus. Finally, we show that pharmacological inhibition of enhancer activity using Minnelide disrupts this chromatin architecture, attenuates the MCL transcriptional program, and exerts antitumor effects against MCL cells *in vitro* and *in vivo*.

## RESULTS

### MCL cells exhibit widespread activation of the enhancer landscape

Large chromosomal aberrations such as the t(11;14) translocation likely exert effects beyond the deregulation of a single oncogene and may reshape higher-order nuclear organization and chromatin states. Nevertheless, these broader regulatory consequences remain insufficiently explored in MCL. To investigate this possibility, we analyzed transcriptional profiles together with chromatin accessibility and the distribution of the activating histone mark H3K27Ac, a marker of active promoters and enhancers, in B cells from MCL patients and control naïve B cells from healthy donors generated within the Blueprint Consortium [17]. Compared to control naïve B cells, MCL cells displayed extensive remodeling of the active chromatin landscape, with more than 1,000 regions showing increased chromatin accessibility and over 11,000 sites with elevated H3K27Ac signal (**Fig. 1a**). Genes located near these regions were enriched for functional categories relevant to lymphoma biology, including B-cell activation, cytokine production, immune signaling, and cell adhesion, as well as processes related to epigenetic regulation such as histone modification (**Fig. 1b**). Consistent with the increased transcriptional activity observed in MCL cells (**Fig. 1d**), most (∼80%) regions with increased H3K27Ac signal were located at gene promoters, while approximately 20% mapped to distal regions likely corresponding to newly activated enhancers (**Fig. 1c**).

**Figure 1.**
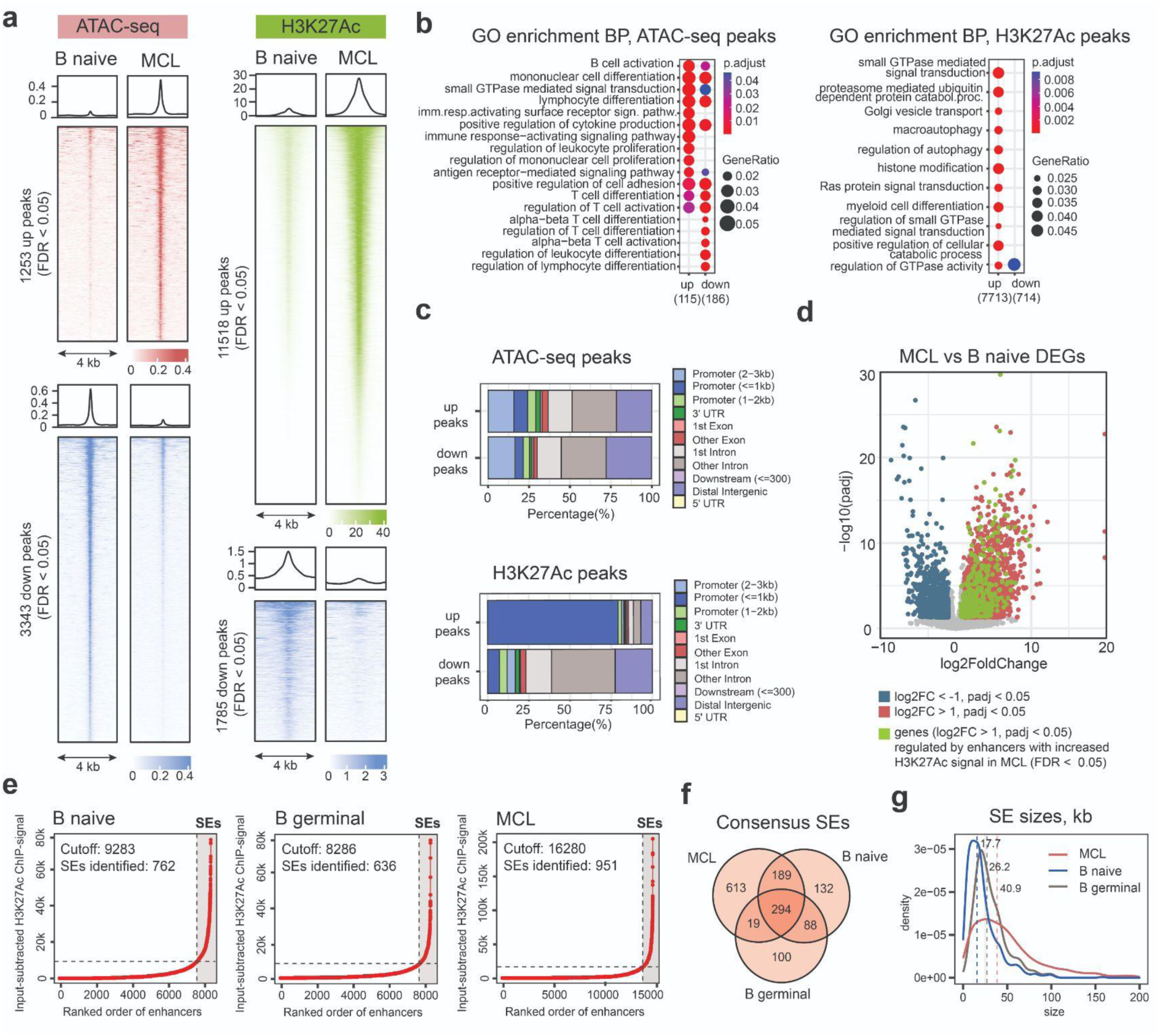
Deregulation of enhancer landscape in MCL cells. **a,** ATAC-seq and H3K27Ac ChIP-seq profiles for differential ATAC and H3K27Ac peaks in MCL patient B cells compared with naïve B cells from healthy donors (FDR < 0.05). **b,** Gene Ontology (GO) enrichment analysis of genes associated with differential ATAC and H3K27Ac peaks (FDR < 0.05). Top 10 enriched biological process categories are shown. **c,** Distributions of the up/down ATAC/H3K27Ac peaks relative to genomic features. **d,** Differentially expressed genes (DEGs) in MCL patient B cells compared with naïve B cells from healthy donors. Upregulated genes (log2FC > 1, padj < 0.05) are shown in red and downregulated genes (log2FC < −1, padj < 0.05) in blue. DEGs predicted to be regulated by enhancers with increased H3K27Ac signal in MCL (FDR < 0.05) are shown in green. Enhancer-gene links were inferred using the ABC model. **e,** Rank ordering of super-enhancers (ROSE) plots for MCL patient samples and control naïve or germinal center B cells (representative biological replicate shown). Each dot represents an enhancer ranked by cumulative H3K27Ac signal. The vertical dashed line indicates the threshold used to define super-enhancers. **f,** Venn diagram showing consensus super-enhancers identified across biological replicates for each condition (MCL, naïve B cells, germinal center B cells) and their overlap between conditions. **g,** Size distributions of consensus super-enhancers in MCL and control naïve or germinal center B cells. Dashed lines indicate median values.

To further characterize enhancer deregulation in MCL, we applied the Activity-By-Contact (ABC) model [18], which integrates ATAC-seq, H3K27Ac ChIP-seq, Hi-C, and gene expression data to infer enhancer-gene interactions (see *Materials and Methods*). ABC analysis in MCL patient samples and control naïve B cells from healthy donors identified 939 enhancers that were newly active or significantly more active in MCL cells, based on increased H3K27Ac signal. These enhancers were predicted to regulate more than 370 genes upregulated in MCL (**Fig. 1d, Table S1**), including the *MYCN* oncogene, *MECOM* and *SRC* proto-oncogenes, and several proliferation and growth-stimulating genes, such as *CCNE1*, *RASGRF1* and *PIK3R2,* supporting a broad association between enhancer activation and transcriptional changes in MCL cells.

Given the extensive enhancer activation observed in MCL, we next examined the landscape of super-enhancers (SEs) - extended clusters of enhancers characterized by exceptionally high levels of transcriptional co-factor binding and strong transcriptional regulatory capacity [19,20]. Using the ROSE algorithm [19] applied to H3K27Ac ChIP-seq data, we identified 1121 super-enhancers active in at least two MCL patient samples, compared with 752 and 520 in naïve and germinal center B cells, respectively (**Fig. 1e**). Approximately 26% (294) of SEs were shared across all three groups (**Fig. 1f**), corresponding to super-enhancers associated with core B-cell identity genes, including MHC II components (e.g. *HLA-DOA, HLA-DPB1*), Ig proteins (*IGL, IGK, IGH*), and the genes of the BCR signaling pathway (e.g. *SYK)* (**Fig. S1a**). Consistent with the presumed origin of MCL from naïve B cells, 17% (189) of MCL SEs were shared exclusively with naïve B cells, whereas only 1.7% (19) overlapped with germinal center B-cell SEs (**Fig. 1f**). Super-enhancers unique to MCL were associated with genes involved in immune signaling and lymphocyte activation, including several genes previously associated with MCL, such as *CD5, CD300A, WEE1,* and *IL2RA*, as well as oncogenic or pro-survival regulators including *PIM1* and *MAP4K1* (**Fig. S1b**). Notably, super-enhancers in MCL were also substantially larger than those in the control B cells, with a median size of ∼40.9 kb compared to 17.7-26.2 kb in naïve and germinal center B cells (**Fig. 1g**). Together, these results indicate extensive remodeling and expansion of the enhancer landscape in MCL cells.

### The translocated *CCND1* locus acquires enhancer-like features and relocates toward the nuclear center in MCL

Given the widespread enhancer activation observed in MCL cells, we next examined whether the t(11;14) breakpoint region itself participates in this enhancer remodeling. Analysis of the derivative chromosome 14 revealed strong enrichment of the activating histone marks H3K27Ac and H3K4me1 across the *IGH* locus (**Fig. 2a**), consistent with the well-established role of *IGH* regulatory elements in driving *CCND1* expression. Although the *IGH* region was classified as a super-enhancer in both control and MCL B cells, the level of H3K27Ac signal was much stronger in MCL samples. Moreover, in MCL cells, these enhancer-associated marks extended beyond the *IGH* locus toward the translocation breakpoint and the *CCND1* region itself, which displayed enhancer-like chromatin features (**Fig. 2a**).

**Figure 2.**
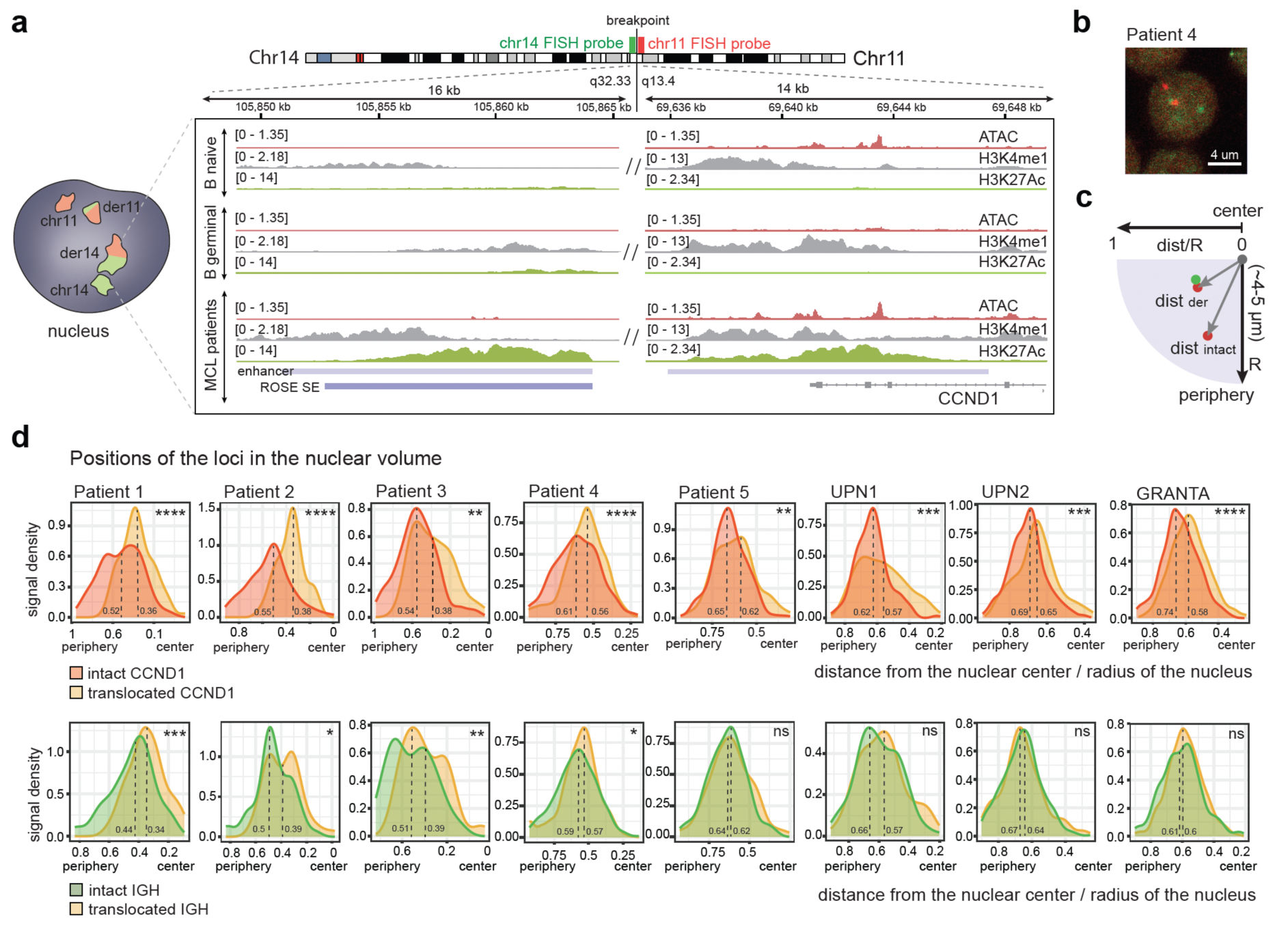
Chromatin state and nuclear positioning of the translocated loci. **a,** ATAC-seq, H3K27Ac, and H3K4me1 ChIP-seq signals at the *IGH* super-enhancer and *CCND1* locus; representative tracks from one of five MCL patients, scaled to one million mapped reads. Blue bars indicate regions called as H3K27Ac peaks (“enhancers”) or ROSE-defined super-enhancers. The translocation breakpoint is located 2.9-220 kb upstream of the *CCND1* locus, with the major breakpoint cluster located ∼109 kb upstream. **b,** Representative confocal image of a B cell carrying the t(11;14) translocation; maximum projection of a z-stack of three 0.5 µm slices. **c,** Schematic representation of the fluorescent probes used to detect the intact *CCND1* and *IGH* loci and the derivative chromosome 14, and the strategy used for signal quantification. For each cell, the distances from intact and translocated signals to the nuclear center were measured and normalized to the nuclear radius, and the resulting values were used to generate population-level distributions of nuclear positioning. **d,** Density plots showing the nuclear distributions of intact (red or green) and translocated (yellow) *CCND1* and *IGH* loci in B cells from five MCL patients and three MCL cell lines (UPN1, UPN2, and GRANTA-519). Statistical significance was determined using the Kolmogorov-Smirnov test for equality of distributions; pval <0.05*, <0.01**, <0.001***, 0.0001****.

As super-enhancers are associated with transcriptionally active nuclear compartments and can influence higher-order chromatin organization, and given that the t(11;14) rearrangement places such a regulatory element at the translocation breakpoint, we next examined whether this structural alteration is accompanied by changes in the nuclear positioning of the affected loci. To address this question, we performed 3D-FISH in B cells from five MCL patients and MCL cell lines using two fluorescent probes. The red probe targeted the region downstream of the translocation breakpoint encompassing the *CCND1* locus, while the green probe targeted the region upstream of the breakpoint and encompassed the locus immediately upstream of the variable *IGH* region. In this configuration, the derivative chromosome 14 could be identified by the overlap of the green and red signals (**Fig. 2b**). The distance of each signal from the nuclear center was measured and normalized to the nuclear radius to obtain the relative spatial distributions of the intact and derivative loci (**Fig. 2c**). We found that the translocated *CCND1* locus consistently occupied a more central nuclear position compared with the intact *CCND1* allele across all MCL patient samples and cell lines examined (**Fig. 2d**). The position of the *IGH* locus, which is normally more centrally located than *CCND1*, either remained unchanged or shifted further toward the nuclear center in some patients. Together, these results indicate that in MCL cells the t(11;14) translocation is associated with the formation of an enhancer-rich regulatory domain around the breakpoint region and *CCND1* locus, accompanied by repositioning of the translocated allele toward the nuclear interior.

### Chromosome 19 is enriched for deregulated genes in MCL

Although *CCND1* overexpression is a hallmark of mantle cell lymphoma, it is not sufficient on its own to drive malignant transformation. In addition to activating *CCND1*, the t(11;14) translocation relocates the *CCND1* locus toward the nuclear center and is associated with the formation of an enhancer-rich regulatory domain around the breakpoint region on derivative chromosome 14. We therefore asked whether genes other than *CCND1* might be affected by this regulatory hub in the MCL context.

To address this question, we analyzed bulk RNA-seq data from nine MCL patients and the GRANTA-519 MCL cell line. Unexpectedly, the largest enrichment of upregulated genes in MCL was not observed on chromosomes 11 or 14. Instead, chromosome 19 emerged as the chromosome most enriched for upregulated genes in MCL patient samples (**Fig. 3a**). When the observed number of upregulated genes was compared with the number expected based on chromosome gene content, chromosome 19 displayed the strongest positive deviation (FDR = 2.2 × 10⁻⁶) (**Fig. 3b**). This pattern was also observed in the GRANTA-519 cell line (**Fig. S2a-b**), and a substantial fraction of chromosome 19 genes upregulated in patient samples overlapped with those upregulated in GRANTA-519 (**Fig. 3c**). Consistent with these observations, positional gene set enrichment analysis (GSEA) revealed a strong enrichment of chromosome 19 genes among the most upregulated transcripts in both MCL patient samples (NES = 2.96, adjusted *p* = 2.3 × 10⁻⁴⁹) and the GRANTA-519 cells (NES = 1.83, adjusted *p* = 1.3 × 10⁻^13^) (**Fig. 3d**).

**Figure 3.**
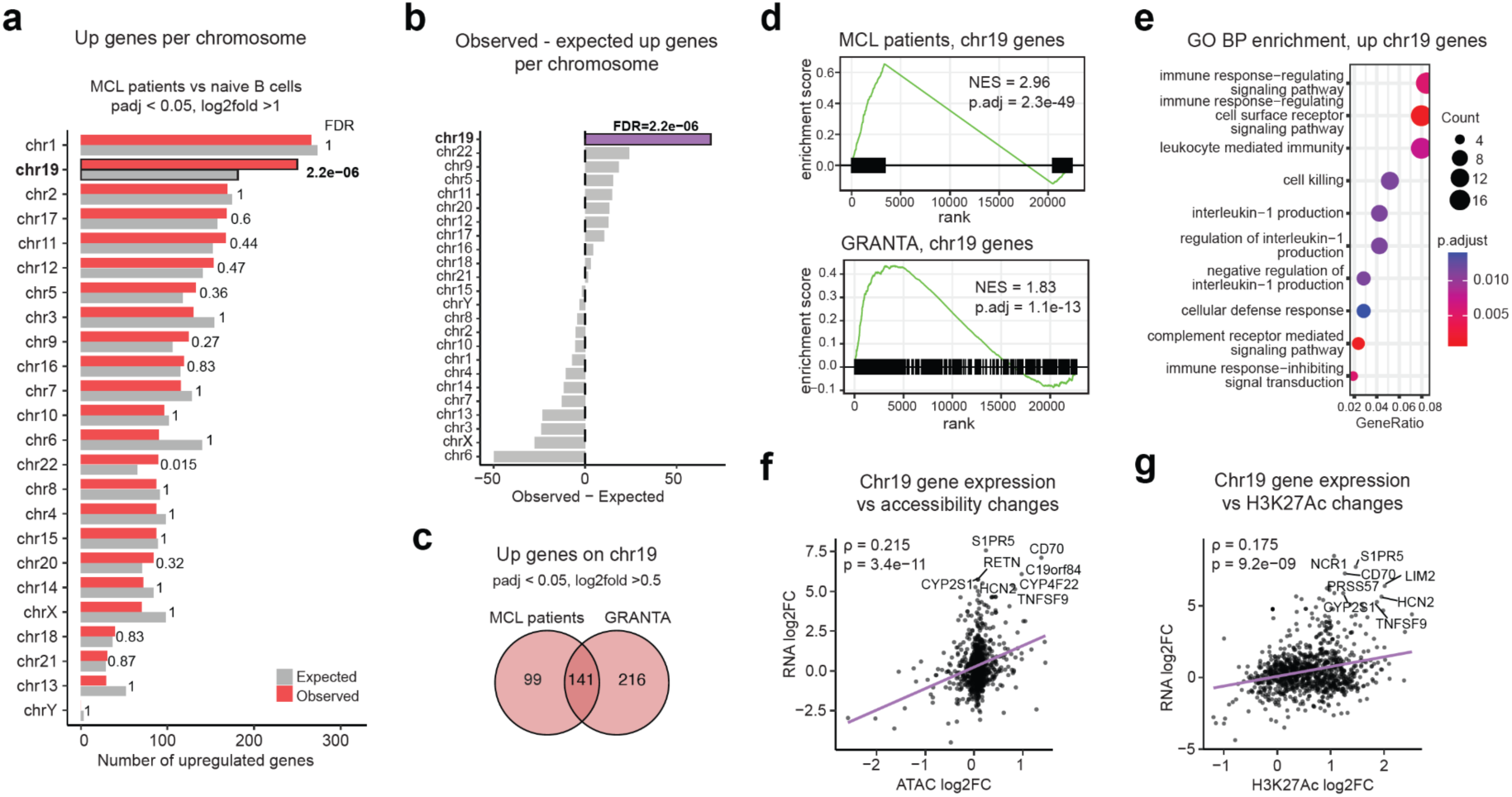
Chromosome 19 is enriched for overexpressed genes in MCL. **a,** Observed vs expected number of upregulated genes in MCL patients compared to control naive B cells (padj <0.01, log2FC > 1). **b,** Difference between the observed and expected numbers of upregulated genes in MCL patients compared to control naive B cells (padj <0.01, log2FC > 1). **c,** Overlap between the upregulated genes on chr19 in MCL patients and GRANTA-519 MCL cell line. **d,** Positional GSEA for the chr19 genes in MCL patients and GRANTA-519 MCL cells. Black bars represent genes ranked according to DESeq2 Wald statistics from the differential expression analysis. Normalized enrichment scores (NES) and Benjamini-Hochberg-adjusted p-values are shown. **e,** Over-representation analysis against the Gene Ontology Biological Process database for the genes upregulated on chr19 in the MCL patients B cells vs control naive B cells (padj <0.01, log2FC > 1). Top 10 enriched categories. **f,g,** Relationship between gene expression changes and chromatin accessibility (f) or H3K27Ac signal (g) changes in MCL patients compared with naïve B cells from healthy donors. Spearman correlation coefficients and corresponding p-values are shown. The top 8 genes ranked by the combined RNA and ATAC/H3K27Ac log2fold change are labeled.

Gene ontology analysis revealed that the upregulated chromosome 19 genes were enriched for immune signaling and leukocyte-mediated immune responses (**Fig. 3e**), functional categories relevant to lymphoma biology. Several of those genes have been previously associated with mantle cell lymphoma, e.g. *BAX, CD70* [21,22], other lymphomas and leukemias, e.g. *BCL3, MAP2K7* [23,24] or have been reported to have a pro-tumorigenic effect on the tumor microenvironment, e.g. *CD70, TYROBP* and *CEACAM1* [22,25–27]. Expression changes of chromosome 19 genes were positively associated with increased chromatin accessibility and H3K27Ac signal at nearby regulatory elements (**Fig. 3f,g**), consistent with a relationship between local chromatin activation and transcriptional upregulation.

Together, these results identify chromosome 19 as a prominent hotspot of transcriptional activation in MCL. These observations prompted us to investigate whether the enrichment of deregulated genes on chromosome 19 might reflect spatial interactions between chromosome 19 and the translocated *CCND1* locus.

### Chr19 interacts with the derivative chromosome 14

Having observed that chromosome 19 is disproportionately enriched for upregulated genes in MCL, we next asked whether this coordinated transcriptional activation might reflect spatial interactions with the translocated *CCND1* locus. Chromosome 19 is known to occupy relatively central positions within the nuclear space [28], similar to other small chromosomes [29,30] and the derivative chromosome 14 carrying the *IGH-CCND1* translocation (**Fig. 2d**). We therefore hypothesized that stochastic interchromosomal contacts between chromosome 19 and the breakpoint-associated regulatory region on derivative chromosome 14 might contribute to the coordinated activation of chromosome 19 genes.

To explore this possibility, we generated Hi-C data from the MCL cell line GRANTA-519 and analyzed published *in situ* Hi-C datasets from five MCL patients and three control naïve B-cell samples [15]. Interchromosomal contacts between chromosome 19 and the chromosome 11 region encompassing the translocation breakpoint were detected in both MCL and control B cells, on both sides of the breakpoint, but with higher contact frequency in MCL cells (**Fig. 4a**). Interestingly, contacts with the breakpoint region were also observed for several other relatively small chromosomes enriched for upregulated genes, such as chromosome 22 (**Fig. 3a-b**; **Fig. 4a**), consistent with the possibility of clustered interchromosomal interactions among these small and mobile chromosomes. Among the detected chr11-chr19 interaction sites, one region on chromosome 19 (site 1) (**Fig. 4a**) coincided with a local peak of transcriptional activation along chromosome 19 (**Fig. 4b**). This site was therefore selected for further validation.

**Figure 4.**
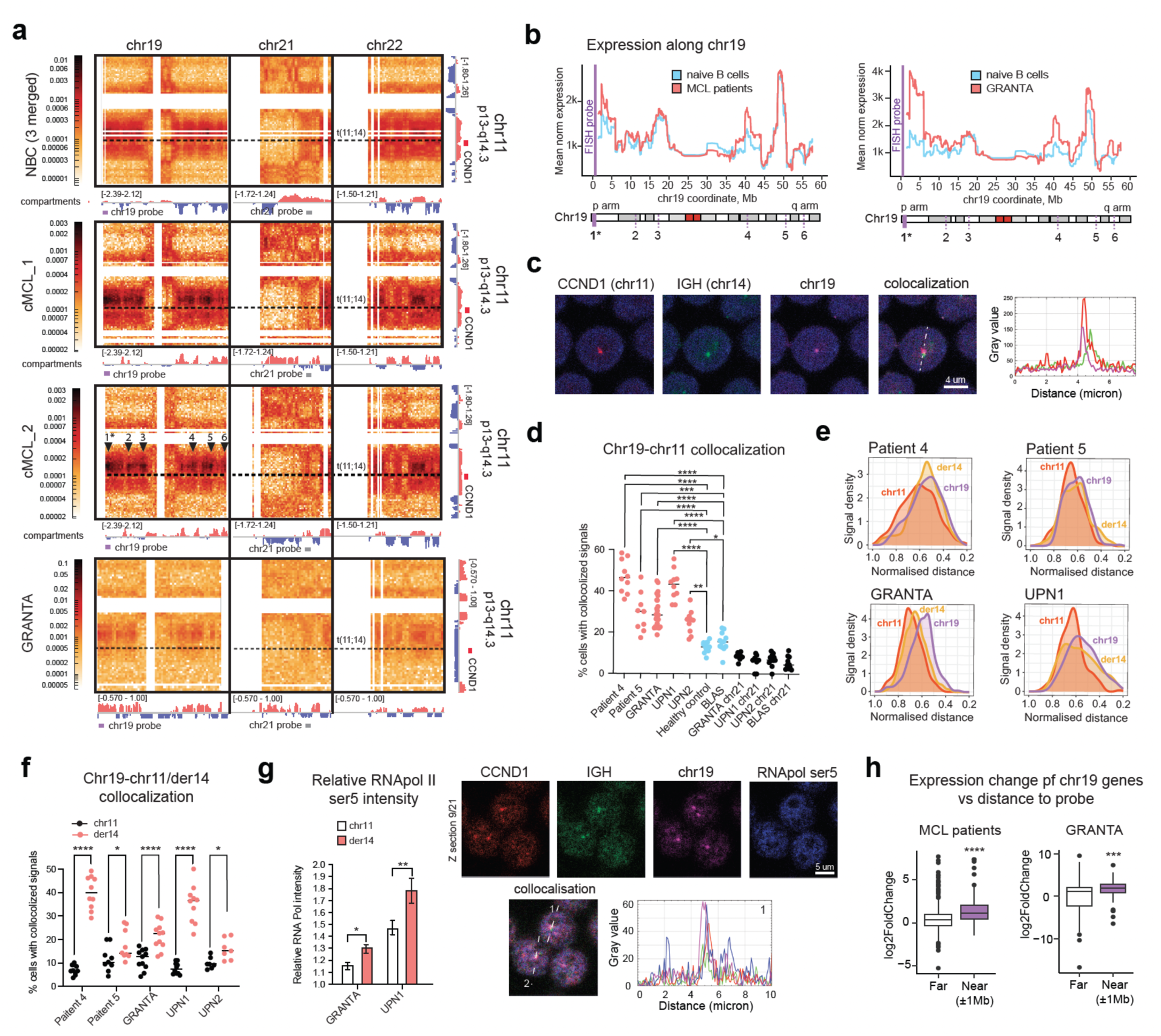
Hi-C and 3D-FISH reveal recurrent interchromosomal contacts between the translocated *CCND1* locus and chr19. **a,** Hi-C maps showing interchromosomal contacts between chr11 breakpoint-adjacent region and the two chromosomes most enriched for upregulated genes in MCL (chromosomes 19 and 22). Chr21, not enriched for upregulated genes, is shown as negative control. Full chromosomes (chr19, chr22) and the chr11 region of interest (p13-q14.3) are shown. Side tracks display eigenvector values assigning active (red) or inactive (blue) compartments. The t(11;14) breakpoint is marked by a dashed line. Arrows indicate contact hotspots; the asterisk marks the chr19 FISH probe region. Positions of chr19 and chr21 FISH probes are schematically indicated. **b,** Mean normalized expression of chr19 genes in MCL patients vs naïve B cells (left) and GRANTA-519 cells vs naïve B cells (right). Values smoothed using a centered rolling mean (window = 101 genes). The vertical line indicates the chr19 FISH probe region. **c,** Representative image of a contact between der14 (red and green) and the chr19 locus (magenta) detected by 3D-FISH in cells from an MCL patient. **d,** Percentage of cells exhibiting a *CCND1*-chr19 contact detected by 3D-FISH in MCL patients, MCL cell lines (UPN1, UPN2, GRANTA-519), a healthy donor control, and a control lymphoblastoid cell line (BLAS). Medians ± SD; each dot represents a technical replicate. Statistical significance determined by one-way ANOVA with Dunnett’s correction. **e,** Density plots of nuclear radial distributions of the intact and translocated *CCND1* loci and the chr19 locus (CTD-2589F14 probe) in B cells from MCL cell lines (GRANTA-519, UPN1) and MCL patients. **f,** Percentage of cells exhibiting contacts between chr19 and the intact or translocated *CCND1* loci in MCL patients and MCL cell lines (UPN1, UPN2, GRANTA-519). Median ± SD; each dot represents a technical replicate. Statistical significance determined by paired t-test. **g,** Left: Intensity of RNA Pol II Ser5 signal within a 1 µm region around the locus, normalized to total nuclear RNA Pol II Ser5 intensity, for intact and derivative chr11 loci in MCL cell lines (UPN1, GRANTA-519). Mean ± SEM. Statistical significance determined by Kruskal-Wallis test with Dunn’s correction. Right: Immuno-3D-FISH with hybridization probes for *CCND1* (red), *IGH* (green), and chr19 (violet), together with antibodies against RNA Pol II Ser5 (blue); representative images and intensity profiles across quadruple colocalization sites are shown. **h,** Expression changes in the MCL vs control comparison for genes located on the chromosome 19p arm either proximal (±1 Mb) or distal to the chr19 probe region. Statistical significance determined by Wilcoxon rank-sum test; horizontal lines represent medians.

To directly test the presence of this interaction, we performed 3D-FISH using probes targeting *CCND1*, the locus immediately upstream of the variable *IGH* region, and the selected chromosome 19 region (chr19:478,637-702,132). In MCL patient samples and cell lines, colocalization between the chromosome 19 probe and the *CCND1* locus was detected in 30-50% of cells (**Fig. 4c,d**). This frequency was substantially higher than the expected probability of random colocalization within the nuclear volume (∼2%) and significantly higher than that observed in control B cells (∼15%) (**Fig. 4d**). As an internal negative control, we quantified contacts between the *CCND1* locus and a region on chromosome 21 lacking detectable Hi-C interactions (**Fig. 4a**); accordingly, chr11-chr21 colocalization remained low in all conditions (**Fig. 4d**). In terms of radial positioning relative to the nuclear center, the translocated *CCND1* locus showed a distribution more similar to chromosome 19 than the intact *CCND1* allele (**Fig. 4e**). Importantly, the interchromosomal interaction occurred predominantly with the derivative chromosome 14 carrying the *IGH-CCND1* translocation rather than the intact *CCND1* allele (**Fig. 4f**), and the 19-*CCND1* contacts frequently colocalized with regions of high RNA polymerase II (Ser5-phosphorylated) signal intensity, with stronger RNA Pol II enrichment observed around the derivative *CCND1* locus (**Fig. 4f,g**).

Finally, we assessed whether transcriptional activation across chromosome 19 was associated with distance from the interaction locus. To avoid confounding effects arising from the centromeric separation between chromosome arms, we restricted the analysis to genes located on the chr19p arm. Genes located within ±1 Mb of the probe region exhibited significantly higher log2 fold-changes than more distal genes in both MCL patient samples (p = 1.7 × 10^-7^) and GRANTA-519 cells (p = 2.3 × 10^-3^) (**Fig. 4h**). Across all chr19p genes, transcriptional activation in MCL patients decreased progressively with increasing genomic distance from the probe region (Spearman ρ = −0.212, p = 1.99 × 10^-7^) (**Fig. S2c**). Linear regression confirmed a significant inverse relationship between distance and expression change (β = −0.358 per log10(bp), p = 6.23 × 10^-10^). To assess whether this pattern was specific to the identified interaction locus, we performed a permutation analysis using 10,000 random probe positions along the chr19p arm. The observed correlation was stronger than that obtained for 99% of randomly sampled loci (empirical p = 0.006) (**Fig. S2d**), indicating that the association is unlikely to arise from an arbitrary genomic position. Similar results were obtained in GRANTA-519 cells (**Fig. S2e-f**).

Together, these results identify recurrent contacts between chromosome 19 and the translocated *CCND1* locus in MCL cells and reveal an association between the interaction site and local patterns of transcriptional activation on chromosome 19.

### Minnelide represses the chromatin landscape of MCL cells and reduces the frequency of chr19-der14 contacts

The results described above revealed an association between transcriptional activation of chromosome 19 genes and spatial interactions involving the *IGH-CCND1* breakpoint region. To test whether perturbing enhancer activity would affect this regulatory architecture, we treated MCL cells with Minnelide, a water-soluble prodrug of triptolide reported to inhibit super-enhancer activity through targeting the XPB subunit of the TFIIH complex [31–33].

We first examined whether Minnelide affects the interchromosomal interaction between chromosome 19 and the derivative chromosome 14. 3D-FISH analysis revealed a significant reduction in chr19-der14 colocalization in both GRANTA-519 and UPN1 MCL cell lines following Minnelide treatment (**Fig. 5a**). This reduction was accompanied by decreased chromatin accessibility both at the chr19 contact sites (**Fig. 5b**) and at the super-enhancer region on derivative chromosome 14 (**Fig. 5c**), together with a modest reduction in *CCND1* expression (**Fig. S3a**).

**Figure 5.**
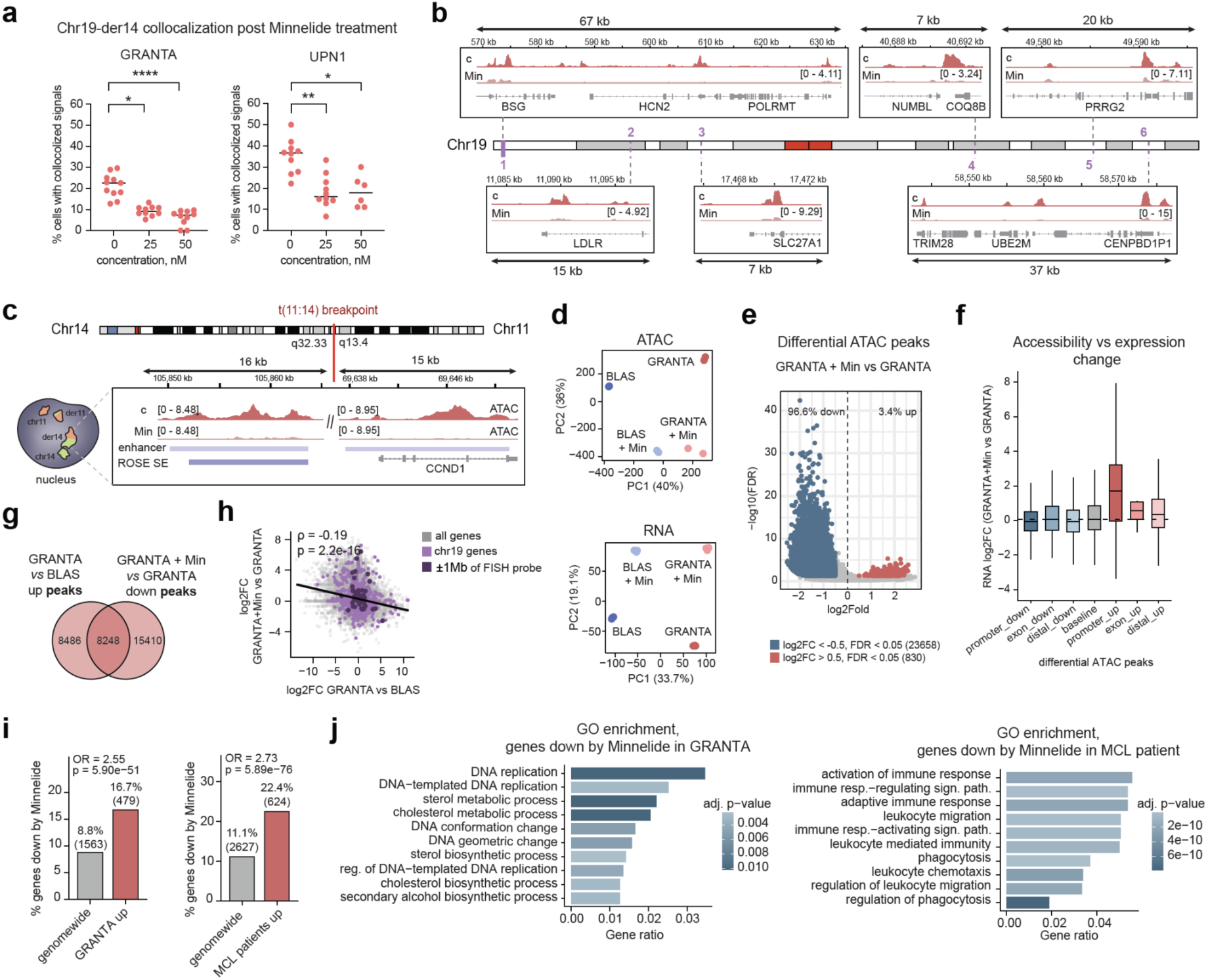
Minnelide perturbs chr19-der14 interactions and partially reverses MCL-associated transcriptional programs. **a,** Percentage of cells exhibiting the chr19-der14 contact before and after Minnelide treatment in the MCL cell lines. **b,** Chromatin accessibility tracks at chr19-chr11 contact sites in GRANTA-519 cells treated or not with 50 nM Minnelide. Signals are scaled to one million mapped reads. **c,** Chromatin accessibility tracks within the super-enhancer (SE) region at the t(11;14) breakpoint in GRANTA-519 cells treated or not with 50 nM Minnelide. Signals scaled to million mapped reads. **d,** Principal component analysis (PCA) of treated and untreated control (BLAS) and MCL (GRANTA-519) cells based on ATAC-seq peaks (top) or gene expression profiles (bottom). **e,** Differential ATAC peaks in GRANTA-519 cells ± Minnelide. Peaks with increased accessibility (log₂FC > 0.5, FDR < 0.05) are shown in red, and peaks with decreased accessibility (log₂FC < −0.5, FDR < 0.05) are shown in blue. **f,** Expression changes following Minnelide treatment for genes associated with ATAC peaks upregulated or downregulated by Minnelide located in promoters, gene bodies, or distal regulatory elements. Horizontal lines indicate medians. **g,** Overlap between ATAC peaks upregulated in GRANTA-519 cells compared with control BLAS cells and peaks downregulated after Minnelide treatment in GRANTA-519 cells. **h,** Relationship between gene expression changes in GRANTA-519 cells vs BLAS controls and expression changes after Minnelide treatment. Spearman correlation coefficients and corresponding p-values are shown. Genes located on chromosome 19 are colored violet, and chr19 genes within ±1 Mb of the chr19 FISH probe are colored dark violet. **i,** Enrichment of transcriptional reversal following Minnelide treatment. Bars show the fraction of genes downregulated by Minnelide among all genes genome-wide (grey) or among genes upregulated in GRANTA-519 cells vs BLAS (left) or in MCL patients vs control naïve B cells (right). Percentages indicate the proportion of Minnelide-downregulated genes, with gene counts shown in parentheses. Enrichment was assessed using Fisher’s exact test; odds ratios (OR) and corresponding p-values are shown. **j,** Gene Ontology enrichment of genes downregulated by Minnelide in GRANTA-519 cells (left) and in MCL patient cells (right). Top 10 enriched categories are shown.

Principal component analysis showed that Minnelide treatment induced a clear shift in both chromatin accessibility and transcriptional profiles genome-wide (**Fig. 5d**). The vast majority (96.6%) of differential ATAC peaks lost accessibility following Minnelide treatment (**Fig. 5e, Fig. S3b**), indicating a general repressive effect of Minnelide on the chromatin state. Both promoter-proximal and distal ATAC peaks were affected, suggesting that Minnelide perturbs chromatin accessibility at both promoters and enhancers (**Fig. S3c**). However, these chromatin changes were not immediately mirrored by corresponding transcriptional changes, as no direct correlation between promoter, enhancer, or gene-body accessibility loss and gene expression reduction was observed (**Fig. 5f**). This may reflect temporal differences between chromatin remodeling and transcriptional responses, although additional time-course experiments would be required to address this possibility.

Of note, Minnelide partially reversed the MCL-associated chromatin activation program: approximately half of the peaks upregulated in GRANTA-519 cells relative to control BLAS cells (8225/16686; 49.3%) were downregulated following Minnelide treatment (**Fig. 5g**). A similar trend was observed at the transcriptional level (**Fig. 5h, Fig. S3d-e**). Expression changes observed in the MCL versus BLAS comparison were negatively correlated with those induced by Minnelide treatment (**Fig. 5h**). This effect was observed genome-wide rather than being restricted to chromosome 19 or any specific loci (**Fig. 5h**), consistent with the broad activity of the drug. Accordingly, genes upregulated in MCL were significantly overrepresented among genes downregulated by Minnelide (OR = 2.55-2.73) (**Fig. 5i**).

Functional analysis revealed that genes downregulated by Minnelide in GRANTA-519 cells were enriched for biological processes related to DNA replication and DNA conformational changes (**Fig. 5j**), consistent with the reported inhibitory effects of triptolide on transcriptional machinery [31,34]. These downregulated genes included chromatin remodelers associated with transcription activation (*CARM1, HMGA1, INO80*), cancer-associated histone methyltransferases (*CARM1, SMYD2, SMYD5, PRMT2, PRMT3*), cell-cycle regulators (*CDK2, FBXO5, CCNE1*) and multiple genes essential for the initiation and maintenance of DNA replication and repair (*MCM3, MCM7, DDX11, POLD2, POLD1, LIG3, NBN, TIMELESS, GINS2, GINS3, RTEL1, CDC45, CDC6, CDC7, TRAIP, DNA2, ZRANB3*). In patient cells, the genes downregulated by Minnelide were enriched for the leukocyte migration and immune-response associated signaling pathways (**Fig. 5j)**. Some examples included the chemokines and their receptors (*CCL5, CCL18, CXCL16, CXCL3, CXCL10, CXCL9, CCR1, CCR2, CCR5, CXCR3, CX3CR1*), growth factors (*PDGFB, VEGFA*), the components and the downstream effectors of the BCR-signaling pathway (*HCK, YES1*, *TXK, SH2D1A*), as well as several known proto-oncogenes (*SRC, KIT, SPI1, YES1*). All of the above-mentioned genes were upregulated in the B cells from MCL patients compared to the naive B cells from healthy donors, and roughly 22% of the overall genes upregulated in MCL were reciprocally downregulated by Minnelide (**Fig. 5i)**.

Together, these results show that Minnelide induces widespread chromatin and transcriptional changes in MCL cells, including reduced accessibility at the breakpoint region, decreased frequency of chr19-der14 contacts, and partial reversal of MCL-associated gene expression patterns. Given the broad activity of the drug, these findings do not establish a specific mechanistic link between disruption of the chr19-der14 interaction and transcriptional changes, but demonstrate that both features are sensitive to pharmacological perturbation.

### Minnelide is effective against MCL cells *in-vitro* and *in vivo*

We next evaluated the effects of Minnelide on MCL cell viability, proliferation, and apoptosis *in vitro*. Minnelide (25-100 nM) reduced the viability of MCL cells as measured by the MTT assay (**Fig. 6a**), with a detectable effect already from day 2 of treatment at 25 nM. Although Minnelide also affected the viability of control lymphoblastoid cell lines derived from healthy donors (BLAS, BLIT, BLYK), the effect was more pronounced and occurred more rapidly in MCL cell lines (GRANTA-519, UPN1, UPN2, Jeko, Mino, NCEB), with the IC₅₀ of 20-25nM in MCL and 40–60 nM in controls (**Fig. S3f**). In contrast, the chronic myelogenous leukemia cell line RPMI8866 was largely unaffected under the conditions tested (**Fig. 6a**). Minnelide treatment induced a dose-dependent decrease in proliferation as measured by the CFSE assay (**Fig. 6b**), with up to ∼80% of cells failing to divide at the highest tested concentration (50 nM), and a similar dose-dependent increase in apoptosis across all tested MCL cell lines (**Fig. 6c**). Depending on the cell line, the proportion of apoptotic cells increased approximately 2-6 fold following treatment with 50 nM Minnelide.

**Figure 6.**
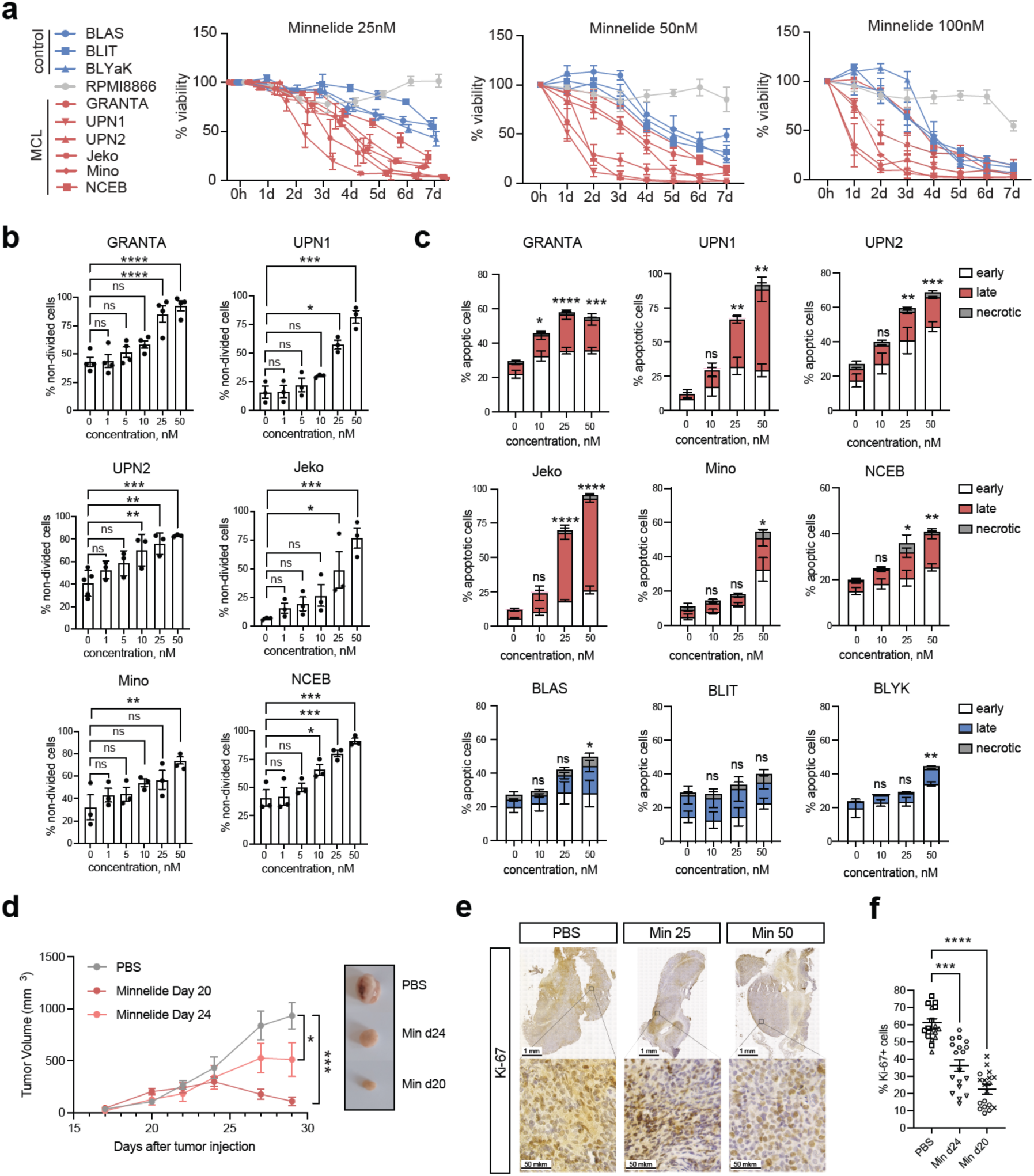
The effect of Minnelide on the viability, apoptosis and proliferation of MCL cells *in vitro* and *in vivo*. **a,** MTT viability assay of MCL and control cells treated with 25-100nM Minnelide for 0-7 days. Mean +/- SEM, N=3-9. **b**, CFSE proliferation assay for the MCL cells treated with 0-50nM of Minnelide for 3 days. The percentage of non-divided cells at the end of the treatment is shown. Mean +/- SEM. Statistical significance determined by one-way ANOVA with Dunnett’s correction. pval <0.05*, <0.01**, <0.001***, 0.0001****, N = 3-5. **c,** Annexin V-PI apoptosis assay for the MCL (red) and control (blue) cells treated with 0-50nM Minnelide for 3 days. Mean ± SEM of apoptotic and necrotic cells. The colors indicate the proportion of early apoptotic (Annexin+, PI-) cells in red, late apoptotic (Annexin+, PI+) cells in blue and necrotic (Annexin-, PI+) cells in gray. Statistical significance determined by one-way ANOVA with Dunnett’s correction. pval <0.05*, <0.01**, <0.001***, 0.0001****, N = 3-6. **d,** Mice tumor volumes on days 15-29 following subcutaneous injections of GRANTA-519 MCL cells. The treatment groups (N = 5) received daily injections of 0.5 mg/kg of Minnelide starting from days 20 or 24 post tumor injection. Control group (N=6) received daily PBS injections starting from day 20. Mean +/- SEM. Statistical significance determined for the values at day 29 using one-way ANOVA with Dunnett’s correction. pval <0.05*, <0.01**, <0.001***. **e,** Representative tumor sections from mice treated or not with Minnelide stained with the anti-Ki-67 antibody. **f,** Percentage of the Ki-67 positive nuclei in the tumor sections from the mice treated or not with Minnelide. Dots represent technical replicates; shapes indicate biological replicates. Mean +/-SEM. Statistical significance determined on technical replicates using Kruskal-Wallis with Dunn’s correction; pval <0.05*, <0.01**, <0.001***, 0.0001****.

We next evaluated the anti-tumor activity of Minnelide *in vivo* using CDX in NSG mice. GRANTA-519 cells were injected subcutaneously into the right flank, and Minnelide treatment was initiated on day 20 or day 24 after injection. In both experimental settings, Minnelide markedly reduced or completely halted tumor growth (**Fig. 6d**). Consistently, immunostaining for the proliferation marker Ki-67 revealed a significant reduction in the proportion of Ki-67-positive cells in xenografts from Minnelide-treated animals (**Fig. 6e,f**).

Together, these results demonstrate that Minnelide exerts a strong anti-tumor effect against mantle cell lymphoma cells both *in vitro* and *in vivo*.

## DISCUSSION

Balanced chromosomal translocations are a hallmark of many lymphoid malignancies and can lead to profound transcriptional deregulation without altering gene dosage. In mantle cell lymphoma, the t(11;14)(q13;q32) translocation drives constitutive *CCND1* expression through enhancer hijacking; however, *CCND1* overexpression alone is insufficient to induce malignant transformation [5,6], indicating that additional regulatory mechanisms contribute to disease pathogenesis.

Our study supports the view that the consequences of t(11;14) in MCL extend beyond local activation of *CCND1* and include broader changes in nuclear organization and chromatin state. We show that the translocated *CCND1* locus repositions toward the nuclear interior, a compartment generally enriched in active chromatin and transcriptional activity [12,35,36]. This observation is consistent with earlier reports showing repositioning of translocated *CCND1* loci in MCL [13,37,38], and fits with the broader principle that spatial repositioning within the nucleus can influence gene regulation [10,11]. In parallel, we detect widespread activation of the enhancer landscape in MCL cells, including increased H3K27Ac signal, expansion of super-enhancer domains, and enhancer-like features extending into the breakpoint-proximal region of the translocated *CCND1* locus. Together, these findings suggest that t(11;14) is associated with the establishment of a more permissive regulatory environment around the translocated allele and genome-wide.

Our results complement a recent work by Oncins et al. [16], showing that t(11;14) in MCL is accompanied by large-scale transcriptional and structural reorganization, including ultra-long-range interactions along the chr11 part downstream of the translocation breakpoint, repositioning of derivative chromosome 11, and alterations of the interchromosomal interaction landscape. Our data add a complementary perspective by focusing on derivative chromosome 14 and by identifying its recurrent interaction with chromosome 19. In this sense, the two studies support a shared model in which translocation-induced effects in MCL are not restricted to the immediate breakpoint environment, but extend across the 3D genome through pre-existing or newly stabilized chromatin contacts.

We identify chromosome 19 as a hotspot of transcriptional activation in MCL. Chromosome 19 is small, gene-dense, and preferentially localized toward the nuclear center ([29], this article), features that may favor interchromosomal interactions. We detect recurrent contacts between chromosome 19 and the *CCND1* locus, preferentially involving the translocated allele, and show that these contacts associate with RNA polymerase II and increased expression of nearby genes. Interestingly, chromosome 22, the second most enriched for upregulated genes in MCL in our data, is also small and centrally positioned and shows Hi-C contacts with the breakpoint region, although we did not proceed with FISH validation. These observations are consistent with the possibility that the nuclear interior provides a permissive environment for interactions between the translocated locus and small, transcriptionally active chromosomes. Although the present study does not establish a direct causal role for these interactions in gene regulation, their recurrence across independent patient samples, cell lines, and orthogonal experimental approaches supports their biological relevance.

Increasing evidence indicates that interchromosomal contacts (ICCs) represent an important structural and regulatory layer of genome organization. Some ICCs are constitutive, such as the nucleolus, where ribosomal genes from multiple acrocentric chromosomes are brought together into a shared transcriptional compartment. Others are more dynamic and stochastic, yet still functionally relevant. For example, olfactory receptor choice depends on stochastic interchromosomal contacts with a trans-acting enhancer [39,40], and IFN-β expression can depend on contacts with several distinct loci that help recruit limiting transcription factors [41]. Co-regulated loci from different chromosomes may also be recruited to shared transcription sites, as reported for globin genes and inducible gene programs in B cells [42–44]. In general, ICCs preferentially involve transcriptionally active and accessible loci rather than silent chromatin [45], and because they are more stochastic than intrachromosomal contacts, they are often detected only in a fraction of cells [46]. This is fully compatible with our observation that chr19-der14 contacts are recurrent but not universal across the cell population.

Our findings further suggest that enhancer-rich chromatin environments may favor such interchromosomal interactions. The breakpoint-associated region displays increased H3K27Ac and accessibility, consistent with a highly active regulatory domain. Enhancer-dense regions and super-enhancers have been linked to long-range interactions and nuclear condensates [47,48], and their deregulation is common across cancers [49–54]. In this context, the chr19-der14 interaction may reflect the combined effects of nuclear positioning, chromatin activity, and regulatory density.

Our data also fit into a broader picture of epigenetic deregulation in MCL. Previous studies have shown that MCL retains substantial epigenetic similarity to naïve mature B cells while simultaneously acquiring tumor-specific changes in chromatin accessibility, compartment status, and DNA methylation [14,15,55]. Disease-associated gains of active compartments and long-range enhancer effects have already been documented in MCL [15,16], and our results add a further layer by linking these regulatory changes to recurrent interchromosomal contacts.

To explore the functional relevance of the chromatin architecture identified here, we perturbed enhancer-associated transcription using Minnelide. Minnelide, a water-soluble derivative of triptolide, is a general transcriptional inhibitor with reported activity against super-enhancer-associated transcriptional programs in cancer cells [33]. Mechanistically, triptolide has been shown to target the CDK7 and XPB subunits of the TFIIH complex [31,56], thereby interfering with transcription initiation. Minnelide has shown antitumor activity in several preclinical cancer models and has advanced to clinical evaluation, including a phase I trial in patients with advanced gastrointestinal cancers [57] and a phase II testing in advanced adenosquamous carcinoma of the pancreas [58].

In our system, Minnelide treatment induced a broad reduction in chromatin accessibility, decreased the frequency of chr19-der14 interactions, and partially reversed the MCL transcriptional program. These observations are consistent with a link between chromatin activity, spatial genome organization, and transcriptional output. At the same time, we acknowledge that the drug does not specifically target the chr19-der14 interaction or the breakpoint-associated regulatory region, and its effects are clearly genome-wide rather than locus-restricted. This lack of specificity limits mechanistic inference: the reduction in interchromosomal contacts could reflect direct perturbation of the relevant regulatory architecture, but it could also arise secondarily from more global transcriptional shutdown and chromatin remodeling. Thus, while our data support the idea that the identified interaction is sensitive to perturbation of enhancer-associated transcription, they do not establish that Minnelide acts primarily through this specific contact. More targeted perturbation strategies will be required to resolve the relative contribution of the breakpoint-associated regulatory domain and the chr19 interaction itself.

The therapeutic results are nevertheless notable. Minnelide showed marked antitumor activity against MCL cells *in vitro* and *in vivo*, consistent with the possibility that MCL cells are particularly dependent on highly active transcriptional and chromatin states. Whether this vulnerability is unique to MCL or extends to other lymphoid malignancies with enhancer-driven oncogenic programs remains an important question. Given the prevalence of enhancer deregulation and transcriptional addiction across cancers, our observations may have broader relevance, although clinical translation will depend on balancing efficacy with the known lack of locus specificity of general transcriptional inhibitors.

In summary, our results indicate that the t(11;14) translocation in MCL is associated with reorganization of nuclear architecture, including recurrent interchromosomal interactions linked to coordinated gene activation. These findings are consistent with a model in which spatial genome organization contributes to disease-specific transcriptional programs and highlight 3D chromatin architecture as an additional layer of regulation in mantle cell lymphoma.

## MATERIALS AND METHODS

### Cells

MCL cell lines GRANTA-519, UPN1, UPN2, Jeko, Mino and NCEB, chronic myelogenous leukemia cell line RPMI8866 and control cell lines BLAS, BLIT and BLYK were used in this study. BLAS, BLIT and BLYK cells were freshly established from the B cells of healthy donors collected in the laboratory with the donors’ informed consent; these cells were immortalized by the EBV (B95-8) transformation and characterized by Genethon (Evry, France). All cells were maintained at 37 °C and 5% CO2 under high humidity and cultured in high-glucose RPMI-1640 medium supplemented with 10% fetal bovine serum (FBS), 2% glucose, 2 mM L-Glutamine, 1 mM Pyruvate, 1000 units/mL of penicillin and 1000 µg/mL of streptomycin (all from Gibco, Thermo Fisher Scientific).

### Patients

Blood samples from hospitalized MCL patients who signed a written informed consent for the study were collected in Gustave Roussy Institute, Villejuif, in accordance with the French legislation. The patient characteristics are described in the table below:

**Table 1.** Characteristics of MCL patients enrolled in the study.

| Patient | Age at blood collection | Sex |
| --- | --- | --- |
| Patient 1 | 69 | Male |
| Patient 2 | 59 | Male |
| Patient 3 | 79 | Male |
| Patient 4 | 73 | Female |
| Patient 5 | 52 | Male |
| Healthy control | 63 | Male |

B cells from patients 1-4 were used in the FISH experiments (see *FISH and ImmunoFISH*) and the RNA-seq experiment without treatments (*see RNA-seq library preparation and sequencing*). B cells from patient 5 were used in the FISH experiments and the RNA-seq experiment with Minnelide treatment (*see RNA-seq library preparation and sequencing*).

### Blood collection and primary B cell isolation

Peripheral blood mononuclear cells (PBMCs) were isolated by Pancoll (PAN Biotech) density gradient centrifugation. PBMC samples from MCL patients in a leukemic phase containing >80% B cells were used directly. Other samples were subjected to B-cell enrichment by negative selection using the MagniSort Human B Cell Enrichment Kit (Thermo Fisher Scientific, #8804-6867-74) prior to downstream analyses.

### FISH probe preparation

The probes for the fluorescent in situ hybridisation (FISH) were prepared from the backmids ordered from BACPAC Genomics. The following backmids were used: RP11-1021J3 (chr11, *CCND1* locus), RP11-346I20 (chr14, upstream of the *IGH* locus), CTD-2589F14 (chr19) and RP11-1056O16 (chr21). The backmids were received as stab cultures, amplified in the DH5α E.coli and purified using the QIAGEN Large-Construct Kit (cat. no. 12462). After purification, the backmids were labeled with fluorophore-dUTPs using the Nick Translation DNA Labeling System 2.0 (Enzo, ENZ-GEN111) according to the manufacturer’s instructions. SEEBRIGHT® Orange 552 dUTPs (ENZ-42842), SEEBRIGHT® Green 496 dUTPs (ENZ-42831) and SEEBRIGHT® Red 650 dUTPs (ENZ-42522) were used to label the chr11, chr14 and chr19/chr21 probes respectively.

### FISH and ImmunoFISH

Cells were resuspended in FBS-free RPMI-1640 at 8 × 10⁶ cells/ml and 80 µl were deposited onto glass coverslips coated with 0.25 mg/ml poly-L-lysine hydrobromide (Merck P1399). After 30 min at 37°C, non-adherent cells were removed with 0.3X PBS. Attached cells were fixed in 4% PFA in 0.3X PBS for 10 min at room temperature, permeabilized in 2% Triton X-100 in PBS for 10 min, and incubated overnight at 4°C in 20% glycerol in PBS. Coverslips were then subjected to three freeze-thaw cycles in liquid nitrogen, treated with 0.1 M HCl for 20 min, and incubated with RNase (200 µg/ml in 2X SSC) for 1 h at 37°C. After equilibration in 50% formamide / 2X SSC, coverslips were stored at 4°C until hybridization.

For each hybridization reaction (two coverslips), nick-translated probes were mixed with 3 µg human COT DNA (Roche 11581074001), ethanol-precipitated, and resuspended in 10 µl in situ hybridization buffer (Empire Genomics). Coverslips were denatured in 70% formamide in 2× SSC and hybridized with the probe mixture after 5 min at 80°C. Hybridization was carried out in a sealed coverslip sandwich at 37°C for 3 days in a humid chamber. After hybridization, coverslips were washed in 2X SSC and PBS. For regular FISH, coverslips were mounted directly. For ImmunoFISH, coverslips were blocked in 0.5% BSA in PBS and incubated with anti-RNA Pol II CTD phospho-Ser5 antibody (Active Motif, 39233; 1:2000) for 2 h at room temperature, followed by Goat anti-Rabbit Alexa Fluor 405 secondary antibody (ThermoFisher, A31556) for 1 h at room temperature. Coverslips were mounted in Vectashield (Eurobio Scientific, H-1000-10). For regular FISH, mounting medium was supplemented with 1.5 µg/ml DAPI (ThermoFisher, D1306); for ImmunoFISH, unstained mounting medium was used. Slides were imaged on a Leica TCS SP8 Multiphoton Confocal Microscope. For each sample, 10 random fields of 291 × 291 µm or 185 × 185 µm were acquired with a Z-step size of 0.5 µm. All images within an experiment were acquired using identical laser intensity, gain, and exposure settings.

### Analysis of FISH images

Three-dimensional images acquired on the Leica TCS SP8 Multiphoton Confocal Microscope were analyzed using Imaris (Bitplane).

For colocalization analysis, FISH signals were detected in each channel using the spot detection function with an estimated spot diameter of 1 µm, followed by manual removal of debris. Only complete cells, defined as cells containing two strong signals for each probe, were retained for analysis. Colocalization between signals was defined using a distance threshold of 1 µm. The percentage of cells exhibiting a given colocalization was calculated relative to the number of complete cells in the field. Analysis was performed separately for 10 fields per experiment, and fields containing ≥50 complete cells were considered technical replicates. Statistical significance was assessed on technical replicates using ordinary one-way ANOVA with Dunnett’s correction, Kruskal-Wallis test with Dunn’s correction, or paired t-test, depending on the comparison (specified in figure legends).

For radial positioning analysis, nuclei were segmented from the DAPI channel and FISH spots were detected as described above. The distance of each spot from the nuclear center was calculated by subtracting the distance to the nuclear membrane from the mean nuclear radius of the field. In experiments analyzing derivative chromosome 14, the distance to the nuclear center was measured relative to the chr11-chr14 colocalized allele. Distances were normalized to the mean nuclear radius of the corresponding field. At least 100 cells were analyzed for patient samples and 200 cells for cell lines. The resulting distributions were visualized as density plots, and differences between distributions were assessed using the Kolmogorov-Smirnov test.

For RNA Pol II colocalization analysis, RNA Pol II signal was segmented from the blue channel and FISH spots were assigned to nuclei. Colocalization was quantified as the ratio of chromosome 19 spot intensity to mean nuclear RNA Pol II intensity. The percentage of colocalizing cells in each field was calculated relative to the number of complete cells and treated as a technical replicate. Statistical significance was evaluated using the Kruskal-Wallis test with Dunn’s correction.

Significance thresholds were defined as p < 0.05 (\**), < 0.01 (*\*\*), < 0.001 (***), < 0.0001 (****).

### MTT cell viability assay

Cells were seeded in 96-well plates (100 µl per well) at 0.5 × 10⁶ cells/ml (GRANTA-519, UPN2, RPMI8866) or 1 × 10⁶ cells/ml (UPN1, Jeko, NCEB, Mino, BLAS, BLIT, BLYK) and treated with 25, 50, or 100 nM Minnelide for 1-7 days in three technical replicates. At the end of treatment, 0.1 mg MTT reagent A (Merck Millipore, CT01-5) was added per well and cells were incubated for 2 h at 37°C. Cells were then lysed overnight in 100 µl MTT lysis buffer (25 mM HCl, 2% acetic acid, 3% DMF, 5% SDS, pH 4.7). Absorbance at 570 nm was measured using a Tecan Infinite F200 PRO plate reader. Background signal from wells without MTT reagent was subtracted. For each condition, the mean of technical replicates was calculated and values were normalized to the untreated control. Experiments were performed in 3–6 biological replicates. Statistical significance was assessed using one-way ANOVA with Dunnett’s multiple comparisons test, comparing treated samples at each time point to untreated controls.

### Annexin V - propidium iodide apoptosis test

0.5 × 10⁶ cells were seeded in 24-well plates and treated with 0, 10, 25, or 50 nM Minnelide for 3 days. After treatment, cells were pelleted and resuspended in 300 µL Annexin buffer (100 mM HEPES pH 7.4, 150 mM NaCl, 2.5 mM CaCl₂). Cells were incubated with 3 µL APC-conjugated Annexin V (BioLegend, #640920) and 3 µL propidium iodide (1 mg/ml) (Sigma-Aldrich, P4170-10MG) for 5 min and analyzed on a BD Accuri C6 Plus flow cytometer (BD Biosciences). Data were analyzed using FlowJo (BD Biosciences) with identical gating applied to all samples within an experiment. Annexin V⁺/PI⁻ cells were classified as early apoptotic, Annexin V⁺/PI⁺ cells as late apoptotic, and Annexin V⁻/PI⁺ cells as necrotic. The percentage of apoptotic cells was calculated as the sum of early apoptotic, late apoptotic, and necrotic cells divided by the total number of gated cells. Experiments were performed in 3-6 biological replicates, and statistical significance was assessed using one-way ANOVA with Dunnett’s multiple comparisons test.

### CFSE proliferation test

Cell proliferation was assessed using carboxyfluorescein diacetate succinimidyl ester (CFSE) staining (CFSE Cell Division Tracker Kit, BioLegend, #423801) according to the manufacturer’s instructions. Briefly, 1 × 10⁶ cells were washed with PBS, resuspended in 1% FBS/PBS at 1 × 10⁶ cells/ml, and labeled with 1 µl of 5 mM CFSE for 4 min at room temperature in the dark. The reaction was stopped by adding 9 ml of 5% FBS/PBS, followed by one wash in 5% FBS/PBS. Cells were then resuspended in growth medium at 5 × 10⁵ cells/ml. Labeled cells were treated with Minnelide (1, 5, 10, 25, or 50 nM) for 3 days. CFSE fluorescence was measured using a BD Accuri C6 Plus flow cytometer (BD Biosciences), and data were analyzed in FlowJo with identical gating applied to all samples within an experiment. The percentage of non-dividing cells (CFSE-positive) was calculated relative to untreated controls. Experiments were performed in 3-5 biological replicates, and statistical significance was assessed using one-way ANOVA with Dunnett’s multiple comparisons test.

### Xenograft studies in NSG mice

GRANTA-519 MCL cells (1 × 10^6^) were suspended in 200 μl of mixture Dulbecco-modified phosphate buffered saline (DPBS, Sigma-Aldrich, Cat#: 59331C) and BD Matrigel Matrix (BD Biosciences; Cat#: 356234) in a ratio of 1∶1, and injected subcutaneously into the right flanks of NSG mice. Tumor growth was monitored every third day using digital calipers. Tumor size was calculated as: Tumor_volume = (length × width^2^)/2. Treatment with Minnelide was started on day 20 or 24 following the injection of the cells. Mice in the treatment groups (N = 5 for each group) received daily intraperitoneal injections of a volume of 0.1 mg/mL Minnelide that achieved a final dose of 0.5 mg/kg of mouse weight. Minnelide was dissolved in PBS. Control animals (N=6) were injected with the equivalent volume of PBS that mirrored the volume of Minnelide per dose. All mice were euthanized on day 29 following the cells injection, tumors were dissected and fixed in 10% formalin for subsequent immunohistochemical analysis.

### Immunohistochemistry analysis

The percentage of Ki-67 positive nuclei was determined using digital images of the tumor section. 9-10 fixed-area hot-spot fields (0,0302 mm²) were quantified within each image. Hot spot fields were defined as regions exhibiting a particularly higher Ki-67 staining compared to surrounding areas. For each hotspot field, the fraction of Ki-67-positive nuclei relative to the total number of nuclei was calculated. The number obtained from one field was considered a technical replica. The fields contained between 1,500 and 2,200 cells in total. Statistical analysis was performed on the technical replicates using the Kruskal-Wallis with Dunn’s correction for multiple comparisons; pval <0.05*, <0.01**, <0.001***, 0.0001****.

### RNA-seq library preparation and sequencing

RNA-seq was performed in two experimental setups. In the first setup, total RNA was extracted from purified B cells from four MCL patients (N = 1 per patient) and the GRANTA-519 MCL cell line (N = 3) without prior treatment. These data were used for comparisons between MCL cells and control B cells from healthy donors (EGAD00001002315) (Figures 1, 3, 4). In the second setup, cells from an MCL patient (patient 5), GRANTA-519, and the control lymphoblastoid cell line BLAS were treated with 50 nM (cell lines) or 25nM Minnelide (patient cells) for 3 days in three technical replicates prior to RNA extraction. These data were used to analyze transcriptional changes following Minnelide treatment (Figure 5). In both setups, total RNA was extracted using the NucleoSpin RNA isolation kit (Macherey-Nagel) according to the manufacturer’s instructions. RNA quantity and quality were assessed using a NanoDrop 2000C spectrophotometer (Thermo Fisher Scientific). cDNA libraries were prepared by Novogene (UK) using the Novogene NGS RNA Library Prep Set (PT042) with poly(A) mRNA enrichment and sequenced on an Illumina NovaSeq 6000 platform in paired-end mode.

### ATAC-seq library preparation and sequencing

Control (BLAS) and MCL (GRANTA-519) cells were treated or not with 50nM Minnelide for 3 days. The ATAC libraries were prepared using the ATAC-Seq Kit (Actif Motif, 53150) according to the manufacturer’s instructions. The libraries were sequenced on the Novaseq6000 platform at the sequencing facility of the Gustave Roussy Institut.

### HiC library preparation and sequencing

Hi-C libraries were prepared as previously described [59]. Briefly, 10⁷ GRANTA-519 MCL cells were crosslinked with 1% formaldehyde for 10 min at room temperature, quenched with 125 mM glycine, washed in PBS, snap-frozen in liquid nitrogen, and stored at −80 °C. After thawing, cells were digested overnight with DpnII (NEB). DNA ends were biotinylated using Klenow enzyme (NEB) with biotin-14-dATP (Invitrogen), followed by ligation with T4 DNA ligase (Thermo Fisher). Crosslinks were reversed at 65 °C with Proteinase K, and DNA was purified by phenol-chloroform extraction, ethanol precipitation, RNase treatment, and cleanup using AMPure XP beads (Beckman Coulter) and Amicon Ultra centrifugal filters (Millipore). DNA was A-tailed with Klenow enzyme, ligation junctions were enriched by biotin pulldown, and Illumina TruSeq adapters were ligated using T4 DNA ligase. Libraries were PCR-amplified and sequenced on an Illumina NovaSeq 6000 platform in paired-end mode.

### RNA-seq analysis

#### Preprocessing

RNA-seq samples from MCL patients and the control (BLAS) and MCL (GRANTA-519) cell lines, treated or not with Minnelide, were prepared and sequenced as described above. In addition, raw transcriptomic data from five additional MCL patients and six naïve B-cell samples from healthy donors were downloaded from EGA (EGAD00001002336, EGAD00001002315) and processed together with the newly generated datasets using an identical computational workflow.

Raw sequencing data were processed using the nf-core/rnaseq pipeline (v3.10.1) [60] with default parameters. Briefly, adapter trimming was performed with TrimGalore (v0.6.7), reads were aligned to the human reference genome GRCh38 using STAR (v2.6.1d), and gene-level counts were quantified with Salmon (v1.9.0). The resulting count matrices were used for downstream analyses in R (v4.3.0).

#### Differential expression analysis

Differential expression analysis was performed with DESeq2 (v1.40.2) [61]. The threshold of adjusted p-value (padj) of less than 0.05 was used to determine statistical significance. The genes with padj < 0.05 and log2fold > 1 were considered upregulated, with padj < 0.05 and log2fold < - 1 were considered downregulated.

#### Chromosomal enrichment analysis

For each chromosome, the observed number of upregulated genes was compared with the expected number calculated as the total number of upregulated genes in the dataset multiplied by the proportion of genes located on that chromosome relative to the total number of analyzed genes. Enrichment significance was evaluated using a one-sided binomial test, and p-values were adjusted for multiple testing using the Benjamini-Hochberg procedure. FDR-adjusted values are reported in the figures.

To assess chromosome-level enrichment across the full ranked transcriptome, positional gene set enrichment analysis (GSEA) using chromosome-specific gene sets was performed. Genes were ranked by the DESeq2 Wald statistic. Chromosome gene sets were generated by grouping annotated genes according to their chromosomal location (chr1-chr22, chrX, chrY). Enrichment analysis was performed with fgsea, using chromosome-specific gene sets containing at least 10 genes. Normalized enrichment scores (NES) and Benjamini-Hochberg-adjusted p-values were calculated for each chromosome.

#### Distance-expression analysis relative to the chr19 FISH probe

Genomic distances between gene loci and the chr19 FISH probe were calculated using GenomicRanges::distanceToNearest function (v1.52.1), and analyses were restricted to genes located on the chr19p arm based on UCSC cytoband annotations (hg38). Distances were log10-transformed prior to analysis. The association between gene expression change (log2FC) and distance from the probe was assessed using Wilcoxon rank-sum test (bar-plots, Figure 4h) and Spearman correlation (scatter plots, Supplementary figure 2c-f). To evaluate whether the observed correlation could arise by chance, a permutation test was performed by sampling 10,000 random probe positions along chr19p and recalculating the correlation. The empirical p-value was defined as the fraction of permutations with absolute correlation coefficients greater than or equal to the observed value.

#### Enrichment of transcriptional reversal by Minnelide

Genes upregulated in MCL cells (GRANTA-519 or patient cells) relative to corresponding controls (BLAS cells or naive B cells from healthy donors) and genes downregulated after Minnelide treatment in MCL cells were identified from differential expression analyses. Enrichment of Minnelide-down genes among MCL-up genes was tested using Fisher’s exact test on a 2×2 contingency table comparing (i) MCL-up vs other genes and (ii) Minnelide-down vs not Minnelide-down. The odds ratio (OR) quantified the enrichment of Minnelide-down genes within the MCL-up gene set relative to the genome-wide background, and significance was assessed using the Fisher exact p-value.

### ATAC-seq analysis

#### Preprocessing

ATAC-seq samples from the control (BLAS) and MCL (GRANTA-519) cell lines were prepared and sequenced as described above. Additional ATAC-seq data from MCL patients and healthy donors were downloaded from EGA (EGAD00001002902, EGAD00001002918). Raw sequencing data were processed using the nf-core/atacseq pipeline (v2.0) [62] with default parameters. Briefly, sequencing adapters were removed with TrimGalore (v0.6.7) and reads were aligned to the human reference genome (GRCh38) using BWA (v0.7.17-r1188).

PCR duplicates were marked using Picard MarkDuplicates (v2.27.4). Alignments were filtered with SAMtools (v1.16.1) to remove duplicates, unmapped reads, secondary alignments, multi-mapping reads, reads mapping to GRCh38 blacklist regions, and mitochondrial DNA, and with BAMtools (v2.5.2) to remove soft-clipped reads, reads with >4 mismatches, and reads with insert size >2 kb. bigWig tracks scaled to one million mapped reads were generated using BEDtools (v2.30.0) and UCSC BedGraphToBigWig (v377) and visualized in IGV. Peak calling was performed using MACS2 (v2.2.7.1) with default parameters. Post-filtering analyses were performed for individual samples and merged replicates, and the merged-replicate tracks are shown in the figures. Profile matrices were generated from merged files using deepTools computeMatrix in reference-point mode.

#### Differential accessibility analysis

The aligned reads and the peaks called by MACS2 separately on each sample were subjected to the differential accessibility analysis in RStudio using DiffBind (v3.10.1) [63]. The peaks with the false discovery rate (FDR) of less than 0.05 were considered differentially accessible. The differentially accessible peaks were annotated to genomic features and the nearest genes using the ChIPseeker R package (v1.36.0) [64].

### ChIP-seq analysis

#### Preprocessing

Datasets from MCL patients and healthy donors were downloaded from EGA (EGAD00001001502, EGAD00001001519, EGAD00001002397). Raw sequencing data were processed using the nf-core/chipseq pipeline (v2.0.0) [65] with default parameters unless stated otherwise. Sequencing adapters were removed using TrimGalore (v0.6.7) and reads were aligned to the human reference genome (GRCh38) with BWA (v0.7.17-r1188). PCR duplicates were marked using Picard MarkDuplicates (v2.27.4). Alignments were filtered with SAMtools (v1.15.1) to remove duplicates, unmapped reads, multi-mapping reads, reads mapping to GRCh38 blacklist regions, and reads mapping to different chromosomes, and with BAMtools (v2.5.2) to remove reads with >4 mismatches or insert size >2 kb. Signal tracks (bigWig) scaled to one million mapped reads were generated using BEDtools (v2.30.0) and UCSC BedGraphToBigWig (v377) and visualized in IGV. Peak calling was performed using MACS2 (v2.2.7.1) with the mappable genome size set to “hs” and q-value cutoff of 0.05. NarrowPeak mode was used for H3K27Ac, and broadPeak mode for H3K4me1. Peaks consistent across replicates or patient samples were identified using ChipR [66] with the minimum number of overlapping peaks set to 2.

#### Differential enrichment analysis

Differential H3K27Ac enrichment analysis was performed in RStudio using DiffBind (v3.10.1) [63] on the aligned reads and the peaks called by MACS2. The peaks with the false discovery rate (FDR) of less than 0.05 were considered differentially enriched. The differentially enriched peaks were annotated to genomic features and the nearest genes using the ChIPseeker R package (v1.36.0) [64].

### HiC analysis

Raw sequencing reads were obtained as described above or downloaded from EGA (EGAD00001006485). Raw fastq files were processed using the nf-core/hic pipeline (v2.1.0) [67] with default parameters and DpnII digestion. Reads were aligned to the human reference genome, filtered to retain valid Hi-C pairs, and converted into multi-resolution contact matrices in mcool format. Contact matrices were normalized using iterative correction and eigenvector decomposition (ICE). The resulting contact maps were visualized using HiGlass [68].

### Identification of enhancers and their targets

Enhancers and their target genes were identified using the activity-by-contact (ABC) model of enhancer-promoter regulation[18], which integrates the ATAC-seq, ChIP-seq, RNA-seq and HiC data. Candidate enhancers were first defined by calling peaks on aligned ATAC-seq data using MACS2. For each enhancer, all expressed genes located within 5 Mb were considered potential targets. For each enhancer-gene pair, the ABC score was calculated as the product of the enhancer activity and the contact frequency between the enhancer and the gene promoter, divided by the sum of the same product calculated for all candidate enhancers within 5 Mb of that gene. Enhancer activity was defined as the geometric fastq mean of the ATAC-seq and H3K27Ac ChIP-seq signals at the element, and contact frequency was derived from Knight-Ruiz normalized Hi-C contact maps. Pairs in which the candidate element overlapped a promoter were excluded. Remaining pairs with ABC score ≥ 0.02 were retained as predicted enhancer-gene interactions. The model was run using the aligned ATAC-seq and H3K27Ac ChIP-seq data described above, DESeq2-normalized RNA-seq counts, and publicly available Hi-C data from the GM12878 lymphoblastoid cell line (ftp://ftp.broadinstitute.org/outgoing/lincRNA/average_hic/average_hic.v2.191020.tar.gz.) Enhancers considered more active in MCL cells were defined as those overlapping regions with significantly increased H3K27Ac signal (FDR < 0.05) by at least 50%.

### Super-enhancer identification

All consistent H3K27Ac peaks passing the MACS2 q-value threshold of 0.05 were considered enhancers. Because promoters can also function as enhancers, TSS-overlapping regions were initially retained. Super-enhancers were identified using the ROSE algorithm [19]. Enhancers located within 12.5 kb were stitched into larger domains to capture dense enhancer clusters. To avoid promoter-driven artifacts, regions within ±2.5 kb of TSSs were excluded during stitching. Stitched and non-stitched enhancers were ranked by input-subtracted H3K27Ac signal, and the slope-1 tangent of the enhancer signal distribution curve was used to define the threshold separating super-enhancers from typical enhancers.

Overlapping super-enhancers were identified using ChIPeakAnno (v3.34.1). Regions with ≥80% overlap across samples within a condition were defined as consensus super-enhancers, and condition-specific super-enhancers were identified using the same overlap threshold. The nearest genomic feature to the center of each super-enhancer was assigned as the associated gene, and enrichment analyses were performed using ChIPeakAnno.

### Concordance between gene expression and chromatin changes

To evaluate the relationship between chromatin accessibility/H3K27Ac signal and gene expression changes, differential ATAC-seq/H3K27Ac ChIP-seq peaks identified by DiffBind were annotated to genes based on genomic proximity with a transcription start site window of ±3 kb using ChIPseeker package (v1.36.0). For each gene, ATAC/H3K27Ac signal changes were aggregated by calculating the mean log2fold change across all associated peaks. RNA-seq log2 fold changes were obtained from the differential expression analysis with DESeq2. Scatter plots were generated to visualize the relationship between aggregated ATAC/H3K27Ac log2fold changes and RNA-seq log2fold changes across genes. The association between chromatin accessibility/H3K27Ac signal and gene expression changes was assessed using Spearman correlation, and statistical significance was evaluated using the corresponding Spearman correlation test p-value.

### Functional enrichment analysis

The functional enrichment analyses were performed on the significantly deregulated genes or genes associated with significantly deregulated ATAC/ChIP peaks distance-wise. Over-representation test (ORA) against the Gene Ontology Database from the clusterProfiler R package (v4.8.2) [69] was used. All the genes detected to be expressed in the MCL and control naive B cells in the RNA-seq experiment were used as the universe. P-values were adjusted using Benjamini-Hochberg correction. Top 10-15 enriched categories by adjusted p-value are shown in the figures.

## Data and code availability

Raw sequencing data from the experiments involving patient material are deposited to EGA and are available on reasonable request. Raw and processed data from the experiments on the cell lines are deposited to an open-access database GEO (GSE291373, GSE291374, GSE291376, GSE291377).

All code used to analyze the data in this study is available on GitHub (https://github.com/annaschwager/mcl_interchrom/).

## Author contributions

Conceptualisation: AS, YV

Methodology: AS, DG, SVU, ET, OD, GA, SVR

Investigation: AS, YG, IT, GR, ET, MP, LL, DAB, SVU

Data analysis: AS, YG, AG, SVU, OD, GA, SVR, DG, YV

Visualization: AS

Clinical samples: VR, JMM

Supervision: YV

Project administration: AS, YV

Writing – original draft: AS

Writing – review & editing: AS, YV

## SUPPLEMENTARY MATERIALS

**Figure S1.**
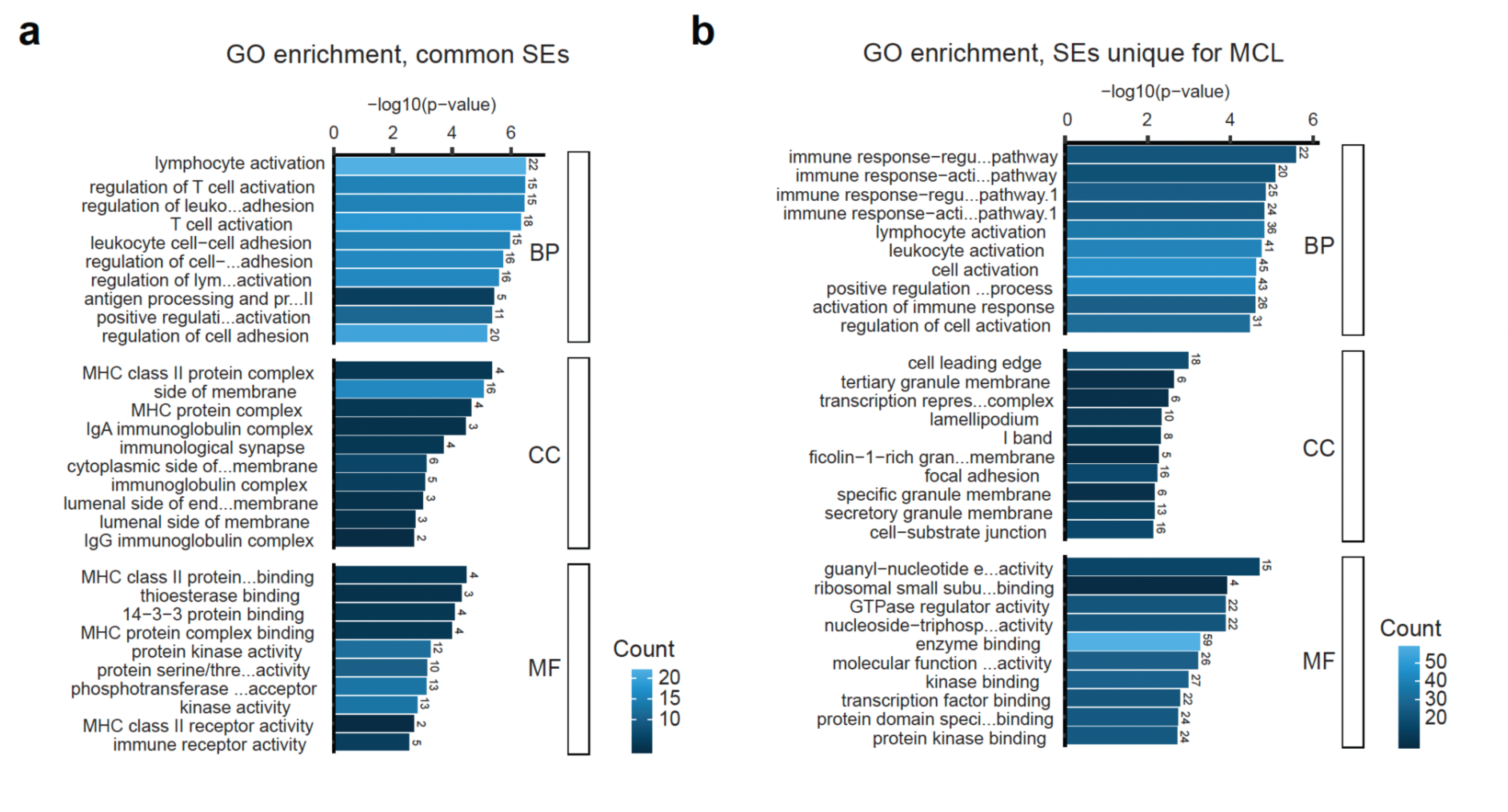
Functional enrichment analysis on super-enhancers. **a,** Gene Ontology enrichment analysis for the genes associated distant-wise with the super-enhancers common for all the three groups of B cells. **b,** Gene Ontology enrichment analysis for the genes associated distant-wise with the super-enhancers unique for MCL.

**Figure S2.**
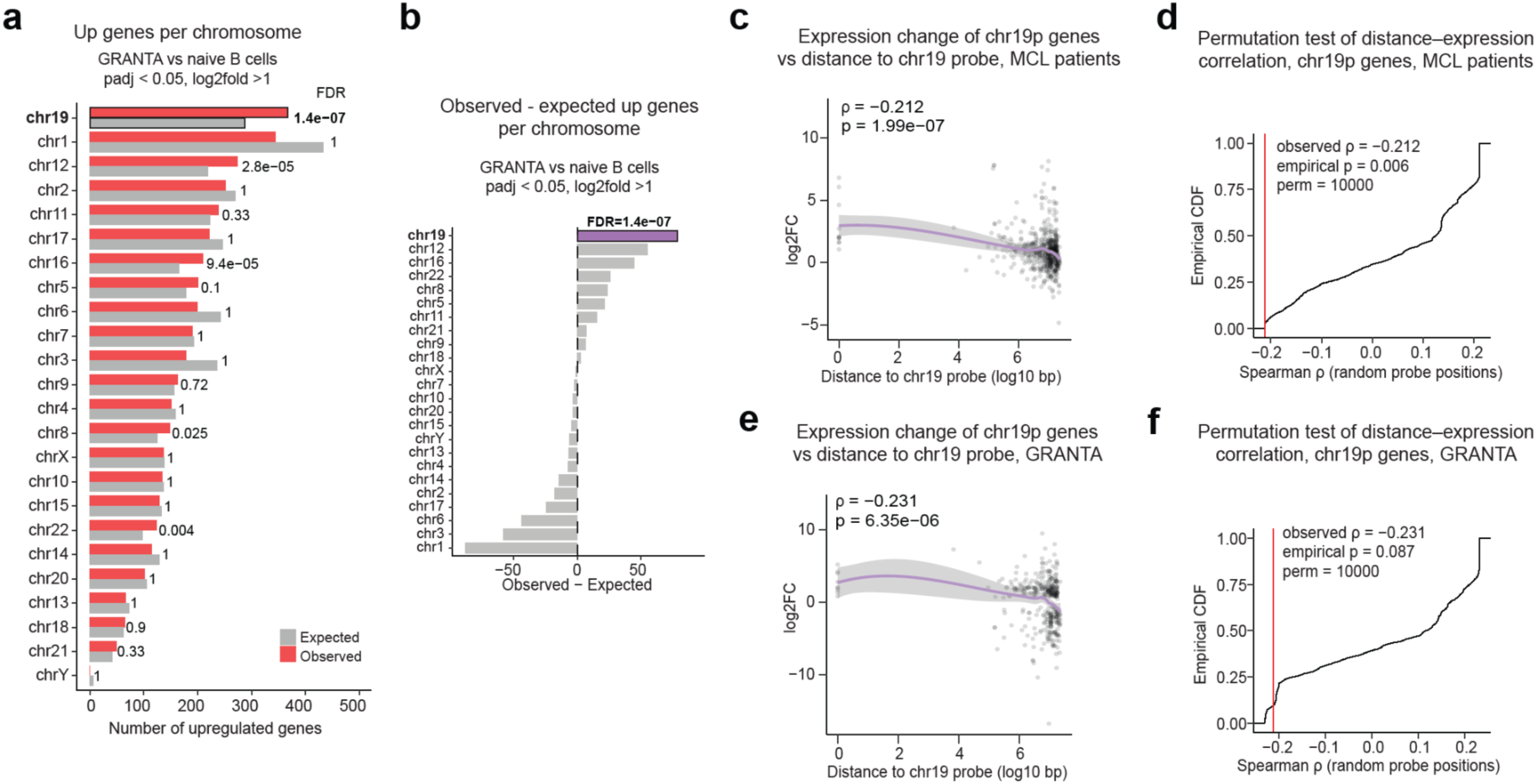
Chr19 gene expression. **a,** Observed vs expected number of upregulated genes in GRANTA-519 cells compared to control naive B cells (padj <0.01, log2FC > 1). **b,** Difference between the observed and expected numbers of upregulated genes in GRANTA-519 cells compared to control naive B cells (padj <0.01, log2FC > 1). **c.** Relationship between gene expression change in MCL patients vs control naive B cells (log2FC) and genomic distance from the chr19 FISH probe for genes located on the chromosome 19p arm. Each point represents a gene; the purple line shows the smoothed trend. Spearman correlation coefficient (ρ) and corresponding p-value are indicated. **d,** Permutation test assessing the significance of the correlation between gene expression change and distance from the chr19 probe. The empirical cumulative distribution of Spearman correlation coefficients obtained from 10,000 random probe positions along chr19p is shown. The red vertical line indicates the observed correlation from panel c; the empirical p-value represents the proportion of permutations with correlation coefficients equal to or more extreme than the observed value. **e,f,** Same as c,d for GRANTA-519 vs control naive B cells comparison.

**Figure S3.**
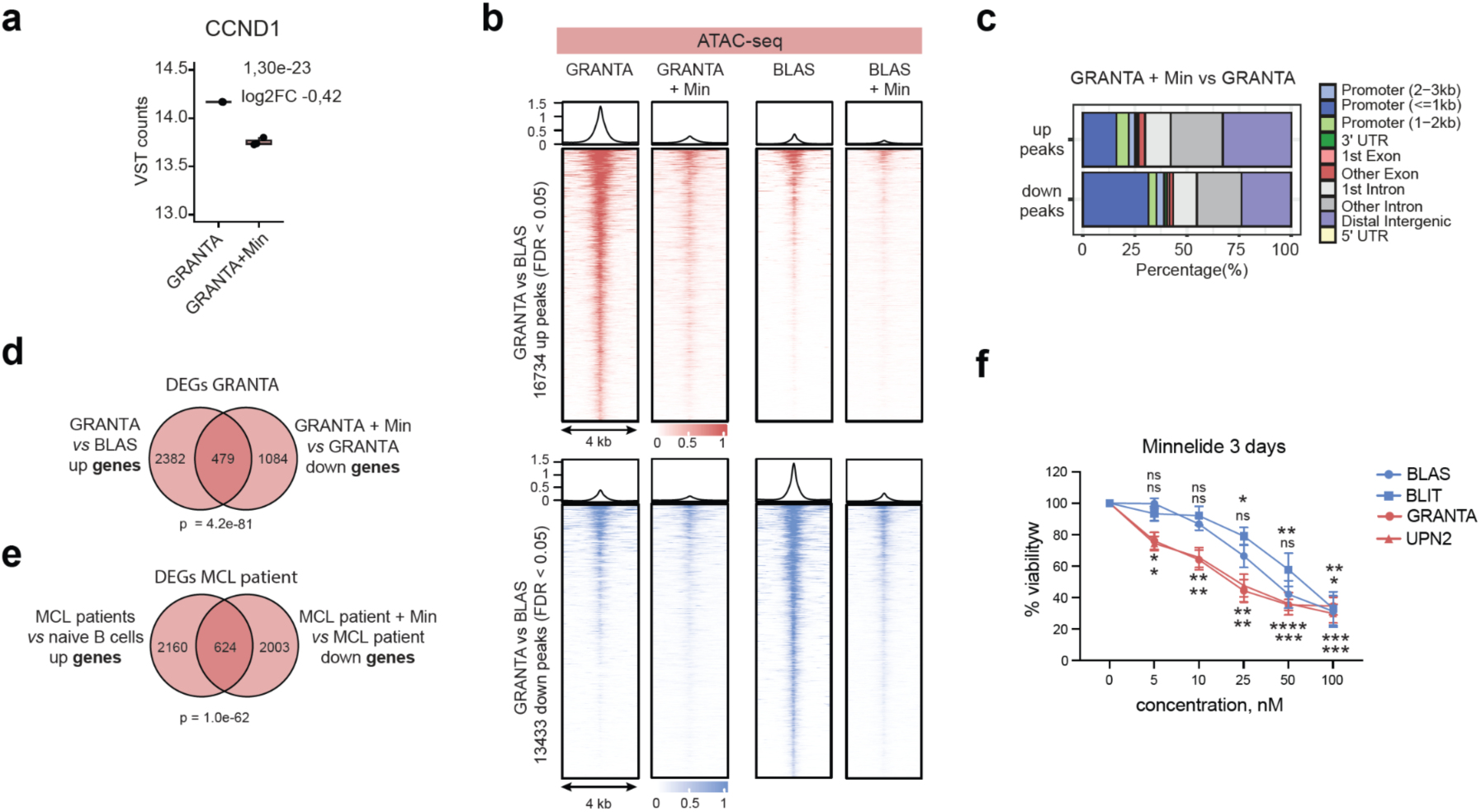
Minnelide treatment. **a**, *CCND1* expression in GRANTA-519 cells following Minnelide treatment. VST-normalized counts are shown. Log2FC and adjusted p-value from the DESeq2 differential expression analysis are indicated. **b**, ATAC-seq profiles of control (BLAS) and MCL (GRANTA-519) cells treated or not with Minnelide at sites of differential accessibility identified in GRANTA-519 cells relative to BLAS controls (FDR < 0.05). **c**, Genomic distribution of ATAC peaks upregulated or downregulated following Minnelide treatment relative to genomic features. **d**, Venn diagram showing the overlap between genes upregulated in GRANTA-519 cells relative to BLAS controls and genes downregulated by Minnelide in GRANTA-519 cells. The p-value from the hypergeometric test is shown. **e**, Venn diagram showing the overlap between genes upregulated in MCL patient cells relative to naïve B cells from healthy donors and genes downregulated by Minnelide in PBMCs from an MCL patient. The p-value from the hypergeometric test is shown. **f,** MTT assay of the viability of MCL (red) and control (blue) cells treated with 0-100nM of Minnelide for 3 days. The data are shown as means +/- SEM, N=3-6. Statistical significance is determined relative to the untreated sample using the two-way ANOVA with Dunett’s correction for multiple comparisons. pval <0.05*, <0.01**, <0.001***, 0.0001****.

## REFERENCES

1. Teras LR, DeSantis CE, Cerhan JR, Morton LM, Jemal A, Flowers CR. 2016 US lymphoid malignancy statistics by World Health Organization subtypes. CA Cancer J Clin. 2016;66:443–59. 10.3322/caac.21357

2. Jain P, Wang M. Mantle cell lymphoma: 2019 update on the diagnosis, pathogenesis, prognostication, and management. Am J Hematol. 2019;94:710–25. 10.1002/ajh.25487

3. Vandenberghe E, De Wolf-Peeters C, van den Oord J, Wlodarska I, Delabie J, Stul M, et al. Translocation (11;14): a cytogenetic anomaly associated with B-cell lymphomas of non-follicle centre cell lineage. J Pathol. 1991;163:13–8. 10.1002/path.1711630104

4. Campo E, Jaffe ES, Cook JR, Quintanilla-Martinez L, Swerdlow SH, Anderson KC, et al. The International Consensus Classification of Mature Lymphoid Neoplasms: a report from the Clinical Advisory Committee. Blood. 2022;140:1229–53. 10.1182/blood.2022015851

5. Fiancette R, Amin R, Truffinet V, Vincent-Fabert C, Cogné N, Cogné M, et al. A myeloma translocation-like model associating CCND1 with the immunoglobulin heavy-chain locus 3’ enhancers does not promote by itself B-cell malignancies. Leuk Res. 2010;34:1043–51. 10.1016/j.leukres.2009.11.017

6. Lovec H, Grzeschiczek A, Kowalski MB, Möröy T. Cyclin D1/bcl-1 cooperates with myc genes in the generation of B-cell lymphoma in transgenic mice. EMBO J. 1994;13:3487–95.

7. Lieberman-Aiden E, van Berkum NL, Williams L, Imakaev M, Ragoczy T, Telling A, et al. Comprehensive mapping of long-range interactions reveals folding principles of the human genome. Science. 2009;326:289–93. 10.1126/science.1181369

8. Dixon JR, Selvaraj S, Yue F, Kim A, Li Y, Shen Y, et al. Topological domains in mammalian genomes identified by analysis of chromatin interactions. Nature. 2012;485:376–80. 10.1038/nature11082

9. Lupiáñez DG, Kraft K, Heinrich V, Krawitz P, Brancati F, Klopocki E, et al. Disruptions of topological chromatin domains cause pathogenic rewiring of gene-enhancer interactions. Cell. 2015;161:1012–25. 10.1016/j.cell.2015.04.004

10. Finlan LE, Sproul D, Thomson I, Boyle S, Kerr E, Perry P, et al. Recruitment to the Nuclear Periphery Can Alter Expression of Genes in Human Cells. PLOS Genetics. Public Library of Science; 2008;4:e1000039. 10.1371/journal.pgen.1000039

11. Kind J, Pagie L, de Vries SS, Nahidiazar L, Dey SS, Bienko M, et al. Genome-wide maps of nuclear lamina interactions in single human cells. Cell. 2015;163:134–47. 10.1016/j.cell.2015.08.040

12. Mota A, Berezicki S, Wernersson E, Harbers L, Li-Wang X, Gradin K, et al. FRET-FISH probes chromatin compaction at individual genomic loci in single cells. Nat Commun. Nature Publishing Group; 2022;13:6680. 10.1038/s41467-022-34183-y

13. Allinne J, Pichugin A, Iarovaia O, Klibi M, Barat A, Zlotek-Zlotkiewicz E, et al. Perinucleolar relocalization and nucleolin as crucial events in the transcriptional activation of key genes in mantle cell lymphoma. Blood. 2014;123:2044–53. 10.1182/blood-2013-06-510511

14. Nadeu F, Martin-Garcia D, Clot G, Díaz-Navarro A, Duran-Ferrer M, Navarro A, et al. Genomic and epigenomic insights into the origin, pathogenesis, and clinical behavior of mantle cell lymphoma subtypes. Blood. 2020;136:1419–32. 10.1182/blood.2020005289

15. Vilarrasa-Blasi R, Soler-Vila P, Verdaguer-Dot N, Russiñol N, Di Stefano M, Chapaprieta V, et al. Dynamics of genome architecture and chromatin function during human B cell differentiation and neoplastic transformation. Nat Commun. Nature Publishing Group; 2021;12:651. 10.1038/s41467-020-20849-y

16. Oncins A, Zaurin R, Toukabri H, Quililan K, Hernández Mora JR, Karpinska MA, et al. Translocations can drive expression changes of multiple genes in regulons covering entire chromosome arms. Nucleic Acids Res. 2025;53:gkaf677. 10.1093/nar/gkaf677

17. Stunnenberg HG, International Human Epigenome Consortium, Hirst M. The International Human Epigenome Consortium: A Blueprint for Scientific Collaboration and Discovery. Cell. 2016;167:1145–9. 10.1016/j.cell.2016.11.007

18. Fulco CP, Nasser J, Jones TR, Munson G, Bergman DT, Subramanian V, et al. Activity-by-contact model of enhancer–promoter regulation from thousands of CRISPR perturbations. Nat Genet. Nature Publishing Group; 2019;51:1664–9. 10.1038/s41588-019-0538-0

19. Whyte WA, Orlando DA, Hnisz D, Abraham BJ, Lin CY, Kagey MH, et al. Master Transcription Factors and Mediator Establish Super-Enhancers at Key Cell Identity Genes. Cell. 2013;153:307–19. 10.1016/j.cell.2013.03.035

20. Lovén J, Hoke HA, Lin CY, Lau A, Orlando DA, Vakoc CR, et al. Selective inhibition of tumor oncogenes by disruption of super-enhancers. Cell. 2013;153:320–34. 10.1016/j.cell.2013.03.036

21. Beltran E, Fresquet V, Martinez-Useros J, Richter-Larrea JA, Sagardoy A, Sesma I, et al. A cyclin-D1 interaction with BAX underlies its oncogenic role and potential as a therapeutic target in mantle cell lymphoma. Proceedings of the National Academy of Sciences. Proceedings of the National Academy of Sciences; 2011;108:12461–6. 10.1073/pnas.1018941108

22. Balsas P, Veloza L, Clot G, Sureda-Gómez M, Rodríguez M-L, Masaoutis C, et al. SOX11, CD70, and Treg cells configure the tumor-immune microenvironment of aggressive mantle cell lymphoma. Blood. 2021;138:2202–15. 10.1182/blood.2020010527

23. Seaton G, Smith H, Brancale A, Westwell AD, Clarkson R. Multifaceted roles for BCL3 in cancer: a proto-oncogene comes of age. Mol Cancer. 2024;23:7. 10.1186/s12943-023-01922-8

24. Chen TJ, Shen Y, Bridges CS, Park CS, Junco JJ, Rabin KR, et al. Pharmacological Inhibition of MAP2K7 Induces Apoptosis and Cell Cycle Arrest in T-Cell Acute Lymphoblastic Leukemia. Blood. 2019;134:3889. 10.1182/blood-2019-129127

25. Xavier S, Nguyen V, Khairnar V, Phan A, Yang L, Nelson MS, et al. CEACAM1 as a mediator of B-cell receptor signaling in mantle cell lymphoma. Nat Commun. 2025;16:4967. 10.1038/s41467-025-60208-3

26. Guo L, An T, Zhou H, Wan Z, Huang Z, Chong T. MMP9 and TYROBP affect the survival of circulating tumor cells in clear cell renal cell carcinoma by adapting to tumor immune microenvironment. Sci Rep. Nature Publishing Group; 2023;13:6982. 10.1038/s41598-023-34317-2

27. Lu J, Peng Y, Huang R, Feng Z, Fan Y, Wang H, et al. Elevated TYROBP expression predicts poor prognosis and high tumor immune infiltration in patients with low-grade glioma. BMC Cancer. 2021;21:723. 10.1186/s12885-021-08456-6

28. Croft JA, Bridger JM, Boyle S, Perry P, Teague P, Bickmore WA. Differences in the localization and morphology of chromosomes in the human nucleus. J Cell Biol. 1999;145:1119–31. 10.1083/jcb.145.6.1119

29. Branco MR, Pombo A. Intermingling of Chromosome Territories in Interphase Suggests Role in Translocations and Transcription-Dependent Associations. PLOS Biology. Public Library of Science; 2006;4:e138. 10.1371/journal.pbio.0040138

30. Bolzer A, Kreth G, Solovei I, Koehler D, Saracoglu K, Fauth C, et al. Three-dimensional maps of all chromosomes in human male fibroblast nuclei and prometaphase rosettes. PLoS Biol. 2005;3:e157. 10.1371/journal.pbio.0030157

31. Titov DV, Gilman B, He Q-L, Bhat S, Low W-K, Dang Y, et al. XPB, a Subunit of TFIIH, Is a Target of the Natural Product Triptolide. Nat Chem Biol. 2011;7:182–8. 10.1038/nchembio.522

32. He Q-L, Titov DV, Li J, Tan M, Ye Z, Zhao Y, et al. Covalent modification of a cysteine residue in the XPB subunit of the general transcription factor TFIIH through single epoxide cleavage of the transcription inhibitor triptolide. Angew Chem Int Ed Engl. 2015;54:1859–63. 10.1002/anie.201408817

33. Noel P, Hussein S, Ng S, Antal CE, Lin W, Rodela E, et al. Triptolide targets super-enhancer networks in pancreatic cancer cells and cancer-associated fibroblasts. Oncogenesis. Nature Publishing Group; 2020;9:1–12. 10.1038/s41389-020-00285-9

34. Vispé S, DeVries L, Créancier L, Besse J, Bréand S, Hobson DJ, et al. Triptolide is an inhibitor of RNA polymerase I and II-dependent transcription leading predominantly to down-regulation of short-lived mRNA. Mol Cancer Ther. 2009;8:2780–90. 10.1158/1535-7163.MCT-09-0549

35. Chen Y, Zhang Y, Wang Y, Zhang L, Brinkman EK, Adam SA, et al. Mapping 3D genome organization relative to nuclear compartments using TSA-Seq as a cytological ruler. Journal of Cell Biology. 2018;217:4025–48. 10.1083/jcb.201807108

36. Su J-H, Zheng P, Kinrot SS, Bintu B, Zhuang X. Genome-Scale Imaging of the 3D Organization and Transcriptional Activity of Chromatin. Cell. 2020;182:1641–1659.e26. 10.1016/j.cell.2020.07.032

37. Sall FB, Pichugin A, Iarovaia O, Barat A, Tsfasman T, Brossas C, et al. Large-scale nuclear remodeling and transcriptional deregulation occur on both derivative chromosomes after Mantle Cell Lymphoma chromosomal translocation [Internet]. bioRxiv; 2020 [cited 2026 Feb 19]. p. 2019.12.30.882407. 10.1101/2019.12.30.882407

38. Schwager AK. Epigenomic and 3D-genomic changes in Mantle Cell Lymphoma [Internet] [thesis]. université Paris-Saclay; 2024 [cited 2026 Feb 19]. https://theses.fr/2024UPASL061. Accessed 19 Feb 2026

39. Lomvardas S, Barnea G, Pisapia DJ, Mendelsohn M, Kirkland J, Axel R. Interchromosomal Interactions and Olfactory Receptor Choice. Cell. 2006;126:403–13. 10.1016/j.cell.2006.06.035

40. Horta A, Monahan K, Bashkirova E, Lomvardas S. Cell type-specific interchromosomal interactions as a mechanism for transcriptional diversity [Internet]. bioRxiv; 2018 [cited 2023 Aug 10]. p. 287532. 10.1101/287532

41. Apostolou E, Thanos D. Virus Infection Induces NF-kappaB-dependent interchromosomal associations mediating monoallelic IFN-beta gene expression. Cell. 2008;134:85–96. 10.1016/j.cell.2008.05.052

42. Osborne CS, Chakalova L, Brown KE, Carter D, Horton A, Debrand E, et al. Active genes dynamically colocalize to shared sites of ongoing transcription. Nat Genet. Nature Publishing Group; 2004;36:1065–71. 10.1038/ng1423

43. Osborne CS, Chakalova L, Mitchell JA, Horton A, Wood AL, Bolland DJ, et al. Myc Dynamically and Preferentially Relocates to a Transcription Factory Occupied by Igh. PLOS Biology. Public Library of Science; 2007;5:e192. 10.1371/journal.pbio.0050192

44. Schoenfelder S, Sexton T, Chakalova L, Cope NF, Horton A, Andrews S, et al. Preferential associations between co-regulated genes reveal a transcriptional interactome in erythroid cells. Nat Genet. Nature Publishing Group; 2010;42:53–61. 10.1038/ng.496

45. Kalhor R, Tjong H, Jayathilaka N, Alber F, Chen L. Genome architectures revealed by tethered chromosome conformation capture and population-based modeling. Nat Biotechnol. Nature Publishing Group; 2012;30:90–8. 10.1038/nbt.2057

46. Noordermeer D, de Wit E, Klous P, van de Werken H, Simonis M, Lopez-Jones M, et al. Variegated gene expression caused by cell-specific long-range DNA interactions. Nat Cell Biol. 2011;13:944–51. 10.1038/ncb2278

47. Rao SSP, Huntley MH, Durand NC, Stamenova EK, Bochkov ID, Robinson JT, et al. A 3D map of the human genome at kilobase resolution reveals principles of chromatin looping. Cell. 2014;159:1665–80. 10.1016/j.cell.2014.11.021

48. Joo J, Cho S, Hong S, Min S, Kim K, Kumar R, et al. Probabilistic establishment of speckle-associated inter-chromosomal interactions. Nucleic Acids Research. 2023;51:5377–95. 10.1093/nar/gkad211

49. Betancur PA, Abraham BJ, Yiu YY, Willingham SB, Khameneh F, Zarnegar M, et al. A CD47-associated super-enhancer links pro-inflammatory signalling to CD47 upregulation in breast cancer. Nat Commun. Nature Publishing Group; 2017;8:14802. 10.1038/ncomms14802

50. Ke L, Zhou H, Wang C, Xiong G, Xiang Y, Ling Y, et al. Nasopharyngeal carcinoma super-enhancer–driven ETV6 correlates with prognosis. Proceedings of the National Academy of Sciences. Proceedings of the National Academy of Sciences; 2017;114:9683–8. 10.1073/pnas.1705236114

51. Wong RWJ, Ngoc PCT, Leong WZ, Yam AWY, Zhang T, Asamitsu K, et al. Enhancer profiling identifies critical cancer genes and characterizes cell identity in adult T-cell leukemia. Blood. 2017;130:2326–38. 10.1182/blood-2017-06-792184

52. Jiang Y-Y, Lin D-C, Mayakonda A, Hazawa M, Ding L-W, Chien W-W, et al. Targeting super-enhancer-associated oncogenes in oesophageal squamous cell carcinoma. Gut. BMJ Publishing Group; 2017;66:1358–68. 10.1136/gutjnl-2016-311818

53. Cohen AJ, Saiakhova A, Corradin O, Luppino JM, Lovrenert K, Bartels CF, et al. Hotspots of aberrant enhancer activity punctuate the colorectal cancer epigenome. Nat Commun. Nature Publishing Group; 2017;8:14400. 10.1038/ncomms14400

54. Yohe ME, Gryder BE, Shern JF, Song YK, Chou H-C, Sindiri S, et al. MEK inhibition induces MYOG and remodels super-enhancers in RAS-driven rhabdomyosarcoma. Sci Transl Med. 2018;10:eaan4470. 10.1126/scitranslmed.aan4470

55. Zhang J, Jima D, Moffitt AB, Liu Q, Czader M, Hsi ED, et al. The genomic landscape of mantle cell lymphoma is related to the epigenetically determined chromatin state of normal B cells. Blood. 2014;123:2988–96. 10.1182/blood-2013-07-517177

56. Manzo SG, Zhou Z-L, Wang Y-Q, Marinello J, He J-X, Li Y-C, et al. Natural product triptolide mediates cancer cell death by triggering CDK7-dependent degradation of RNA polymerase II. Cancer Res. 2012;72:5363–73. 10.1158/0008-5472.CAN-12-1006

57. Borazanci E, Saluja A, Gockerman J, Velagapudi M, Korn R, Von Hoff D, et al. First-in-Human Phase I Study of Minnelide in Patients With Advanced Gastrointestinal Cancers: Safety, Pharmacokinetics, Pharmacodynamics, and Antitumor Activity. The Oncologist. 2024;29:132–41. 10.1093/oncolo/oyad278

58. Skorupan N, Ahmad MI, Steinberg SM, Trepel JB, Cridebring D, Han H, et al. A phase II trial of the super-enhancer inhibitor Minnelide^TM^ in advanced refractory adenosquamous carcinoma of the pancreas. Future Oncol. 2022;18:2475–81. 10.2217/fon-2021-1609

59. Zakharova VV, Magnitov MD, Del Maestro L, Ulianov SV, Glentis A, Uyanik B, et al. SETDB1 fuels the lung cancer phenotype by modulating epigenome, 3D genome organization and chromatin mechanical properties. Nucleic Acids Res. 2022;50:4389–413. 10.1093/nar/gkac234

60. Patel H, Manning J, Ewels P, Garcia MU, Peltzer A, Hammarén R, et al. nf-core/rnaseq: nf-core/rnaseq v3.23.0 - Gallium Gecko [Internet]. Zenodo; 2026 [cited 2026 Mar 21]. 10.5281/zenodo.18805594

61. Love MI, Huber W, Anders S. Moderated estimation of fold change and dispersion for RNA-seq data with DESeq2. Genome Biol. 2014;15:550. 10.1186/s13059-014-0550-8

62. Patel H, Espinosa-Carrasco J, Langer B, Ewels P, bot nf-core, Garcia MU, et al. nf-core/atacseq: [2.1.2] - 2022-08-07 [Internet]. Zenodo; 2023 [cited 2026 Mar 21]. 10.5281/zenodo.8222875

63. Stark R, Brown G. DiffBind: Differential binding analysis of ChIP-Seq peak data. :75.

64. Yu G, Wang L-G, He Q-Y. ChIPseeker: an R/Bioconductor package for ChIP peak annotation, comparison and visualization. Bioinformatics. 2015;31:2382–3. 10.1093/bioinformatics/btv145

65. Patel H, Espinosa-Carrasco J, Wang C, Ewels P, bot nf-core, Silva TC, et al. nf-core/chipseq: nf-core/chipseq v2.1.0 - Platinum Willow Sparrow [Internet]. Zenodo; 2024 [cited 2026 Mar 21]. 10.5281/zenodo.13899404

66. Newell R, Pienaar R, Balderson B, Piper M, Essebier A, Bodén M. ChIP-R: Assembling reproducible sets of ChIP-seq and ATAC-seq peaks from multiple replicates. Genomics. 2021;113:1855–66. 10.1016/j.ygeno.2021.04.026

67. Servant N, bot nf-core, Ewels P, Garcia MU, Talbot A, Peltzer A, et al. nf-core/hic: nf-core/hic v2.1.0 [Internet]. Zenodo; 2023 [cited 2024 June 21]. 10.5281/zenodo.7994878

68. Kerpedjiev P, Abdennur N, Lekschas F, McCallum C, Dinkla K, Strobelt H, et al. HiGlass: web-based visual exploration and analysis of genome interaction maps. Genome Biology. 2018;19:125. 10.1186/s13059-018-1486-1

69. Wu T, Hu E, Xu S, Chen M, Guo P, Dai Z, et al. clusterProfiler 4.0: A universal enrichment tool for interpreting omics data. The Innovation. 2021;2:100141. 10.1016/j.xinn.2021.100141

